# Multiple distinct metastatic cell states are induced by epithelial-mesenchymal plasticity

**DOI:** 10.1101/2025.08.15.670583

**Authors:** Fahda Alsharief, Robert K. Suter, Apsra Nasir, Idalia Cruz, Gray W. Pearson

**Affiliations:** Lombardi Comprehensive Cancer Center and Department of Oncology, Georgetown University, Washington, DC 20057, USA; Department of Laboratory Sciences, College of Applied Medical Sciences, Jouf University, Sakaka, Saudi Arabia

**Author notes:** Equal contribution (Fahda Alsharief, Robert K. Suter). To whom correspondence should be addressed. Gray W. Pearson Lombardi Comprehensive Cancer Center Georgetown University 3970 Reservoir Road NW, Washington, DC 20057.

## Abstract

Epithelial-mesenchymal transition (EMT) enables epithelial cancer cells to acquire mesenchymal-associated traits that can promote invasion and metastasis. Although distinct EMT-associated states have been linked to invasive and metastatic behavior, it remains unclear when these states arise during primary tumor progression, how they diversify, and whether metastatic competence is restricted to a particular EMT phenotype. Using single-cell RNA sequencing in a genetically engineered mouse model of triple-negative breast cancer (TNBC), together with functional studies of tumor organoids, we reconstructed the emergence of EMT-associated heterogeneity during tumor progression. We found that early malignant cells first lost mammary lineage identity, generating lineage-altered epithelial states with increased intrinsic plasticity. Rather than progressing through a single EMT program, these plastic states diversified through ERK1/2-low and ERK1/2-high EMT-associated programs. These programs generated distinct hybrid epithelial-mesenchymal states in early tumors and more uniform mesenchymal-like subpopulations at later stages, with canonical EMT features, diminished plasticity, and highly invasive behavior. Importantly, metastatic competence was not restricted to a single EMT-associated state—both heterogeneous hybrid cells and more uniform mesenchymal-like cells initiated metastases, with metastatic lesions retaining features of their initiating populations. Together, our results show that EMT-associated heterogeneity in TNBC emerges through early lineage-state disruption followed by parallel regulatory programs that generate distinct metastatic cell states rather than converging on a single highly metastatic phenotype.

## INTRODUCTION

The epithelial–mesenchymal transition (EMT) is a cellular plasticity program in which epithelial cells lose features such as apical–basal polarity and junctional adhesion while acquiring mesenchymal-associated properties, including cytoskeletal remodeling, extracellular matrix remodeling, and increased motility (1). EMT is essential during embryonic development and is transiently reactivated during tissue repair (2). In carcinomas, however, aberrant activation of EMT-associated programs can promote invasion, dissemination, therapeutic resistance, and metastatic progression (3, 4). EMT is not a binary conversion from a fully epithelial to a fully mesenchymal state. Instead, cancer cells can occupy a spectrum of epithelial, hybrid epithelial-mesenchymal, and mesenchymal-like states (5, 6). This epithelial-mesenchymal plasticity can generate substantial phenotypic heterogeneity within tumors (7), but how EMT-associated heterogeneity emerges during tumor progression remains incompletely understood.

EMT is regulated by signaling networks that integrate extrinsic cues from the tumor microenvironment with intrinsic transcriptional circuitry (8). Although EMT has often been framed around a conserved set of EMT-inducing transcription factors, including SNAI1/2, TWIST1, and ZEB1/2, it is increasingly clear that EMT regulatory mechanisms are highly context dependent (9, 10). Distinct extracellular cues, including TGFβ family ligands, receptor tyrosine kinase ligands, extracellular matrix changes, biomechanical forces, inflammatory signals, and metabolic stresses, can activate different signaling pathways and produce EMT-associated states with distinct molecular and phenotypic properties (11–13). These states may differ in the extent of epithelial marker loss, mesenchymal gene induction, transcription factor usage, reversibility, and invasive behavior (14–18). As a result, standard EMT markers such as Vimentin expression or loss of E-Cadherin may not fully capture the diversity or functional significance of EMT-associated tumor cell states (19). A major unresolved challenge is to define how EMT-associated heterogeneity is generated within primary tumors, where multiple signals, regulatory mechanisms, and tumor cell states coexist.

The functional consequences of EMT-associated heterogeneity are also incompletely understood. EMT can promote invasion by increasing motility, extracellular matrix remodeling, and transitions between collective and single-cell invasion (20–23). However, EMT-associated states can also differ in proliferation, survival, plasticity, and dependence on specific microenvironmental conditions (24–27). Hybrid epithelial-mesenchymal states may retain epithelial traits that support growth or collective behavior while also acquiring invasive properties, whereas more mesenchymal-like states may display enhanced motility but altered fitness or state stability (5, 21, 28–30). Studies in multiple tumor models suggest that metastatic ability is not always restricted to the most mesenchymal cells (5, 26, 27). However, how distinct EMT-associated states arise during primary tumor progression and how these states contribute to metastatic behavior remain poorly defined.

Triple-negative breast cancer (TNBC) provides an important setting in which to address these questions. TNBCs lack expression of estrogen receptor, progesterone receptor, and HER2, limiting the targeted treatment strategies available for other breast cancer subtypes (31). Clinically, TNBCs are associated with aggressive disease, early recurrence, and increased risk of metastasis (32). At the molecular and cellular levels, TNBCs frequently display basal-like or claudin-low features, elevated EMT-associated programs, and substantial intratumoral heterogeneity, including epithelial, hybrid, and mesenchymal-like tumor cell populations (33–35). Although EMT-associated plasticity has been strongly implicated in the invasive, metastatic, and therapy-resistant behavior of TNBC (21, 36–38), the mechanisms that generate EMT heterogeneity, the timing of its emergence during tumor evolution, and the EMT-associated states that support metastatic competence remain poorly defined.

Defining how EMT-associated heterogeneity arises requires experimental systems that can resolve heterogeneous tumor cell states across stages of tumor progression. Human tumors provide essential clinical relevance, but clinical samples are usually collected at limited time points and cannot readily reconstruct the temporal sequence through which EMT-associated states emerge and diversify (39, 40). Genetically engineered mouse models provide an opportunity to examine tumor evolution over time, but breast cancer models differ substantially in the extent and character of EMT-associated phenotypes they produce (5, 41–43). Models that capture a broader epithelial, hybrid, and mesenchymal-like spectrum are therefore needed to define how EMT-associated heterogeneity emerges and how distinct states relate to metastatic behavior. To address this problem, we used the C3(1)-TAg model, a basal-like TNBC genetically engineered mouse model that develops spontaneous mammary tumors with extensive EMT-associated heterogeneity (21, 22, 41, 44, 45).

Here, we used single-cell RNA sequencing in the C3(1)-TAg model, together with functional studies of tumor-derived organoids, to define how EMT-associated heterogeneity emerges during TNBC progression. This approach allowed us to reconstruct the timing, regulatory organization, and functional consequences of EMT-associated state diversification within primary tumors. Our findings support a model in which EMT-associated heterogeneity arises through early disruption of epithelial lineage identity followed by diversification into distinct regulatory and phenotypic states with metastatic capacity. These results provide a framework for understanding how EMT generates multiple metastasis-competent states in TNBC that produce distinct metastatic phenotypes rather than a single uniform metastatic endpoint.

## RESULTS

### C3-TAg tumor progression generates extensive EMT-associated transcriptional heterogeneity

To define the organization of epithelial–mesenchymal transition (EMT)-associated heterogeneity during tumor progression, we performed single-cell RNA sequencing (scRNA-seq) on the C3(1)/SV40 T-antigen (C3-TAg) mouse model of triple-negative breast cancer (TNBC). This model develops spontaneous ductal carcinoma in situ–like lesions that progress to invasive and metastatic tumors with EMT heterogeneity similar to that observed in human TNBC (22, 38, 45). We analyzed normal mammary epithelium, 5 mm tumors representing early invasive lesions, and >15-mm tumors representing advanced disease **(Fig. 1A)**. After quality-control filtering and doublet exclusion, 2,561 high-quality cells were retained for analysis **(Fig. S1A)**. Unsupervised clustering of the full dataset identified 16 transcriptionally distinct cell populations, which were annotated using canonical lineage markers, transcriptional similarity to a reference mouse mammary cell atlas (46), cell-cycle status, and sample origin **(Fig. S1A–I)**. UMAP visualization resolved the expected epithelial, stromal, hematopoietic, endothelial, smooth-muscle, and neuronal populations of the mammary gland and tumor microenvironment **(Fig. S1C–E)**. Within the malignant compartment, C3-TAg tumor epithelial cells occupied two visually distinct UMAP lobes: one enriched for 5-mm tumor cells and a second composed almost exclusively of cells from >15-mm tumors **(Fig. S1A, C)**. Each lobe contained multiple clusters, indicating progressive diversification of tumor epithelial states with tumor progression **(Fig. S1B)**.

**Figure 1.**
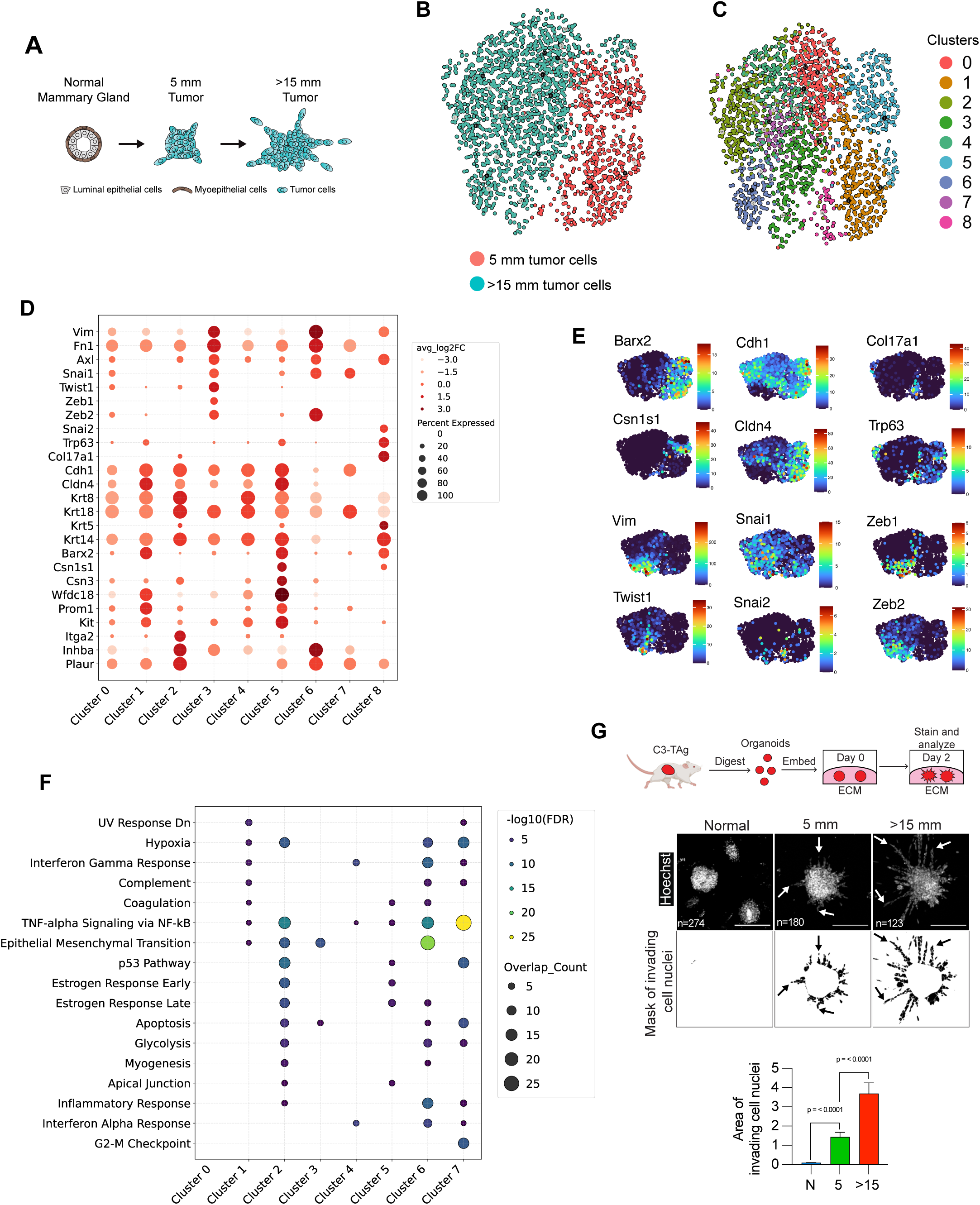
C3-TAg tumor progression generates extensive EMT-associated transcriptional heterogeneity. **A.** Graphical model showing the conditions analyzed by scRNA-seq. n=3 samples/condition. **B.** UMAP showing annotation of 5 mm and >15 mm tumor cells after clustering when controlling for the contribution of cell cycle genes and when just plotting tumor cells. **C.** UMAP showing Seurat clusters from 5 mm and >15 mm tumor cells. **D.** Bubble plot showing the expression of the indicated genes across the tumor cell clusters. **E.** UMAPs show the expression of the indicated genes in tumor cells. **F.** Bubble plot showing the enrichment of Hallmark Pathway genes in the tumor cell clusters. **G.** Invasion of organoids derived from normal mammary gland, 5 mm and >15 mm tumors.

To focus on EMT-induced heterogeneity rather than transient differences in cell cycle phase, we removed canonical cell-cycle genes and re-clustered tumor-derived epithelial cells **(Fig. 1B)**. This analysis partitioned tumor epithelial cells into nine transcriptional clusters **(Fig. 1C, S1J, Table S1)**. Cells from 5 mm tumors were largely confined to Clusters 1, 5, and 8, indicating limited transcriptional diversity at this early invasive stage **(Fig. 1B–C)**. Although these cells expressed Krt14, atlas-based correlation showed that they were more closely related to luminal progenitor cells than to basal or myoepithelial lineages **(Fig. 1D, S1I, K)**. This interpretation was supported by elevated expression of luminal progenitor markers, including Prom1, Kit, and Barx2 (47, 48), and lactation-associated genes, including Wfdc18, Csn1s1, and Csn3 (46) **(Fig. 1D–E, S1K)**. Cluster 8 exhibited elevated p63 and basal-associated features, including Krt5 and Col17a1 (49), suggesting lineage infidelity; however, this population did not expand in >15 mm tumors, indicating that luminal-to-basal conversion is not a major driver of progression in this model **(Fig. 1D–E, S1J)**. This lineage relationship mirrors human TNBCs, which are transcriptionally closest to luminal progenitor cells despite frequent basal keratin expression (49, 50). In contrast to 5 mm tumors, cells from >15 mm tumors were distributed across eight of the nine clusters, reflecting broad phenotypic diversification **(Fig. 1B–C)**.

Analysis of hallmark gene sets (51) identified EMT as one of the most strongly enriched programs among cluster-defining genes, with the highest enrichment in Cluster 6 and additional enrichment in Clusters 2, 3, and 8 **(Fig. 1F, Table S2)**. EMT enrichment was markedly more prominent in >15-mm tumors than in 5-mm tumors, both because advanced tumors contained a larger fraction of cells in EMT-enriched clusters and because these cells occupied multiple distinct EMT-associated states. Consistent with this shift, >15-mm tumor cells showed reduced at-las-based similarity to luminal progenitor cells and increased similarity to mesenchymal stem–like populations **(Fig. S1L)**. Organoids derived from >15-mm tumors also showed increased invasive behavior compared with organoids from 5-mm tumors (Fig. 1G), linking transcriptional diversification and EMT to functional heterogeneity.

The EMT-enriched clusters differed in epithelial feature loss, mesenchymal gene induction, and EMT transcription factor expression. Cluster 8 resembled a p63-positive hybrid EMT state previously defined by our group (14, 52). Cluster 2 represented a distinct hybrid state marked by Itga2, Inhba, Spp1, and Plaur expression. Cells in this cluster also retained epithelial character and showed limited expression of the EMT mesenchymal marker Vimentin and canonical EMT transcription factor (Snai1, Snai2, Twist1, Zeb1, Zeb2) expression (53) **(Fig. 1D–F)**. In contrast, Clusters 3 and 6 showed more classical EMT features, including broader expression of Vimentin, Fn1, and Axl (22, 54, 55) and reduced expression of cell–cell adhesion genes including Cdh1/E-Cadherin and Cldn4 (56, 57). The two clusters were distinguished by their EMT transcription factor and ECM regulatory gene profiles **(Fig. 1D–F, Table S3)**. Thus, C3-TAg tumor progression is marked by a transition from epithelial-like states with limited transcriptional heterogeneity to advanced tumors containing multiple hybrid and mesenchymal-like EMT-associated states.

### Trajectory analysis identifies divergent ERK1/2-low and ERK1/2-high EMT programs

To define relationships among EMT-associated states, we used Monocle3 to infer transcriptional trajectories among tumor epithelial cells (58). We interpreted pseudotime within the biological framework provided by our sampling of 5-mm and >15-mm tumors, allowing transcriptional progression to be evaluated in relation to tumor evolution. Trajectory analysis resolved a bifurcating structure terminating in Clusters 3 and 6 rather than a single linear epithelial-to-mesenchymal continuum **(Fig. 2A–C, S2A)**. Notably, p63-expressing Cluster 8 cells did not align with the main trajectory trunk or either EMT branch, but instead mapped as a separate endpoint of phenotypic variation **(Fig. 2C, S2A)**. This organization is consistent with our previous work defining p63-positive cells as a stable hybrid EMT state (14, 52) and suggests that Cluster 8 is not an obligatory intermediate in progression toward Clusters 3 or 6.

**Figure 2.**
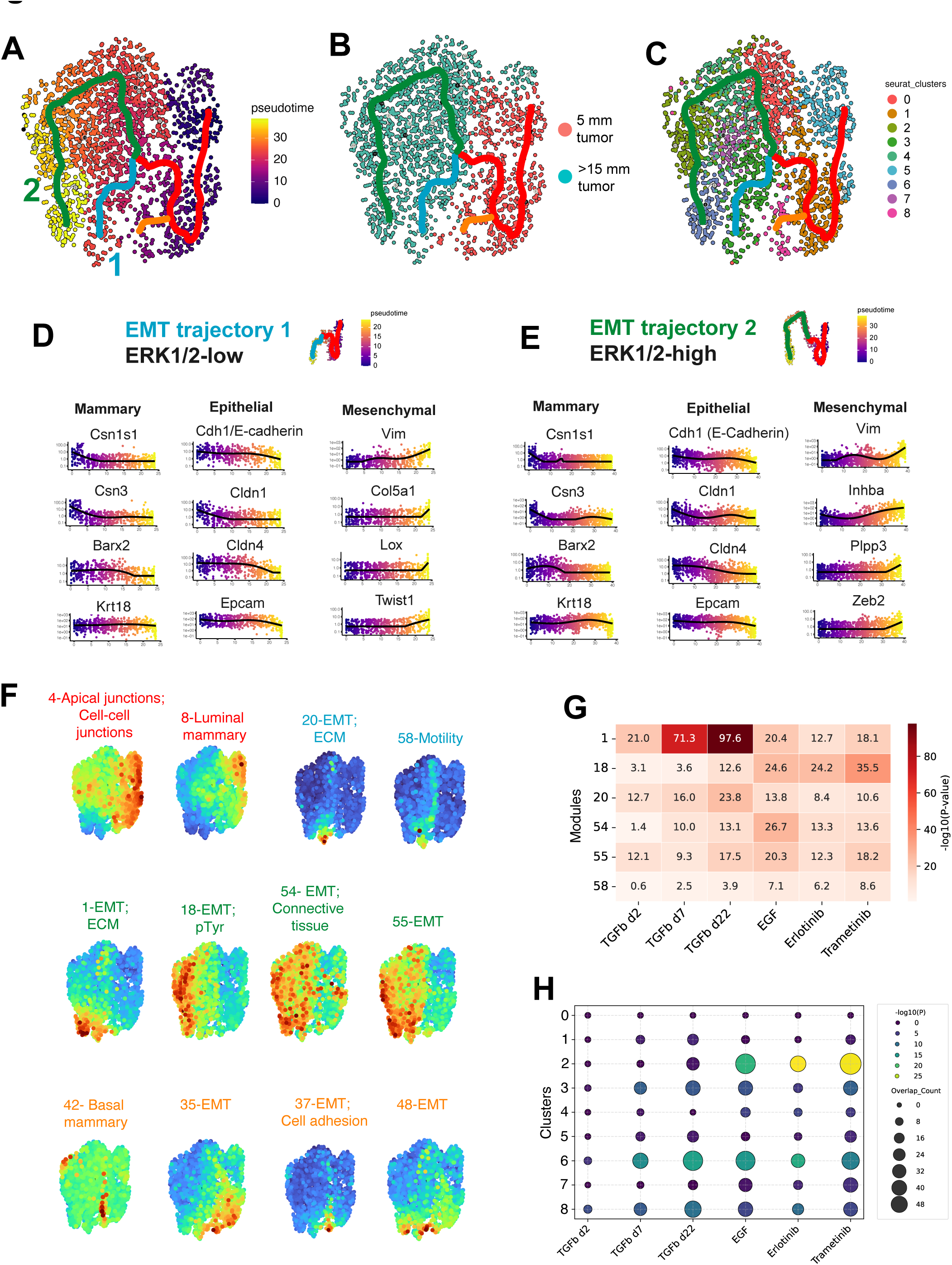
EMT-associated tumor cell heterogeneity emerges along two transcriptional trajectories during C3-TAg tumor progression. **A.** Monocle3 trajectory analysis of tumor cells from 5-mm and >15-mm C3-TAg tumors. UMAP shows cells colored by pseudotime. **B.** Monocle3 trajectory analysis of tumor cells from 5-mm and >15-mm C3-TAg tumors. UMAP shows cells colored by tumor stage. **C.** Monocle3 trajectory analysis of tumor cells from 5-mm and >15-mm C3-TAg tumors. UMAP shows cells colored by Seurat cluster identity. **D.** Expression of representative mammary lineage, epithelial, and mesenchymal-associated genes across pseudotime for EMT trajectory 1. **E.** Expression of representative mammary lineage, epithelial, and mesenchymal-associated genes across pseudotime for EMT trajectory 2. **F.** UMAPs show the indicated gene-module activity scores. **G.** Heatmap shows enrichment analysis comparing the indicated gene module signatures with EMT programs induced TGFβ or EGFR-ERK1/2 signaling in tumor organoids. **H.** Dot plot showing enrichment analysis comparing Seurat cluster gene signatures with EMT programs induced by TGFβ or EGFR-ERK1/2 signaling in tumor organoids.

Pseudotemporal analysis revealed shared early events across both EMT branches. Among the top 100 differentially expressed genes, both trajectories first showed downregulation of mammary lineage–defining genes, including Barx2, Wfdc18, Csn1s1, and Csn3, together with reduced expression of cell–cell junction components such as Cldn1 and Cldn4 **(Fig. 2D–E)**. These early changes occurred during the transition from 5 mm to >15 mm tumors and preceded broad mesenchymal gene induction. After branching, both trajectories showed progressive induction of Vim and Zeb2 and coordinated reduction of the canonical epithelial markers Epcam and Cdh1/E-Cadherin **(Fig. 2D–E)**. However, the two branches differed in EMT transcription factor composition—Trajectory 1 showed increased Twist1, Zeb1, and Prrx1 expression, whereas Trajectory 2 cells preferentially upregulated Sox5 **(Fig. 2D–E, S2B–C)**. The EMT branches were also distinguished by differences in the specific mesenchymal-associated genes represented among the top differentially expressed genes **(Fig. 2D–E, S2B–C)**. Thus, C3-TAg tumor cells diversify into parallel EMT-associated trajectories with shared early lineage and epithelial feature loss but distinct mesenchymal transcriptional program activation.

Gene-module analysis further supported this organization **(Fig. S2D, Table S4)**. Modules associated with mammary identity and cell-cell junctions declined along both trajectories **(Fig. 2F, S2D–E, Tables 5–9)**. In contrast, the branches diverged in modules associated with EMT, motility, and ECM reorganization **(Fig. 2F, S2D–E, Tables S5-9)**. Because we previously found that TGFβ and EGFR-ERK1/2 pathway activation induce distinct EMT-associated programs in C3-TAg organoids (22), we next tested whether trajectory-associated modules were enriched for genes regulated by these pathways. Modules active at the endpoints of both trajectories, as well as genes enriched in terminal Clusters 3 and 6, overlapped with TGFβ-induced programs, particularly genes activated after prolonged TGFβ exposure **(Fig. 2F–G, Table S10)**. Thus, both trajectories engage programs that can be stimulated by a canonical EMT-inducing signal. In contrast, Trajectory 2-aligned modules were specifically enriched for ERK1/2-regulated genes, including Itga2, Inhba, Lama3, and the autocrine EGFR ligands Areg and Ereg **(Fig. 2E–G, S2F, Table S10)**. Genes highly expressed in Cluster 2, an intermediate state along Trajectory 2, and Cluster 6, the Trajectory 2 endpoint, were also enriched for ERK1/2-regulated genes **(Fig. 2H, Table S11).** Based on these features, we refer to Trajectory 1 as the ERK1/2-low EMT trajectory and Trajectory 2 as the ERK1/2-high EMT trajectory.

Together, these results indicate that EMT activation in C3-TAg tumors diverges into at least two programs that share early loss of mammary lineage and epithelial identity but differ in downstream regulatory composition, including differential engagement of ERK1/2-regulated genes.

### EMT-associated plasticity is acquired during tumor progression and generates stable mesenchymal-like states

Pseudotime predicts the likely order of transcriptional changes during EMT but cannot determine whether tumor cells functionally undergo these transitions or whether the resulting states remain reversible. Because EpCAM declined progressively along both EMT trajectories and can be measured on live cells, we used EpCAM expression to isolate EpCAM-high hybrid-like cells and EpCAM-low mesenchymal-like cells from >15 mm tumors. Tumor cells were separated by fluorescence-activated cell sorting (FACS) into EpCAM-high (E-hi) and EpCAM-low (E-lo) fractions, cultured as organoids and subsequently reanalyzed by EpCAM-based FACS analysis to assess transition between the two states **(Fig. 3A).** E-hi organoids consistently generated E-lo progeny, whereas E-lo organoids rarely regained high EpCAM expression **(Fig. 3B)**. Consistent with this directional transition, unsorted primary tumor organoids became progressively enriched for E-lo cells over time **(Fig. S3A)**. Moreover, new tumors initiated from E-lo organoids remained composed predominantly of E-lo cells, indicating that the mesenchymal-like state is stably inherited during both in vitro organoid culture and in vivo orthotopic tumor growth **(Fig. S3B)**.

**Figure 3.**
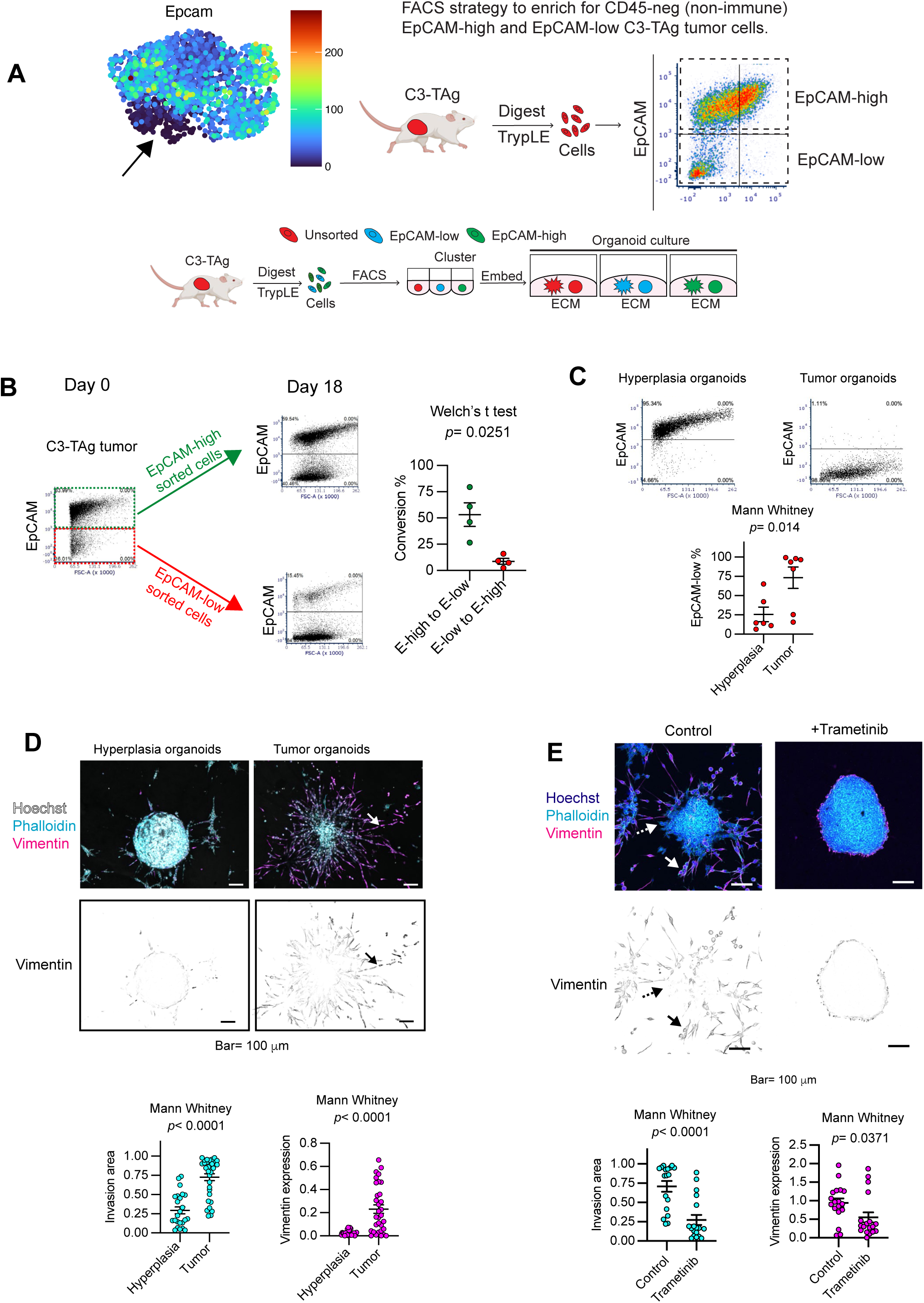
EMT-associated plasticity is acquired during tumor progression and generates stable mesenchymal-like states. **A.** Experimental workflow for isolating CD45-negative C3-TAg tumor cells by FACS based on EpCAM expression. EpCAM-high and EpCAM-low tumor cell fractions were sorted and cultured as tumor organoids to assess epithelial plasticity and invasive behavior. UMAP feature plot shows Epcam expression across tumor cells from the scRNA-seq dataset. **B.** Representative FACS plots and quantification of plasticity between EpCAM states after organoid culture. **C.** Representative FACS plots and quantification comparing EpCAM-low cell abundance in organoids derived from hyperplastic lesions and tumors. **D.** Representative immunofluorescence images and quantification of low- and high-plasticity tumor organoids stained for Hoechst, phalloidin, and Vimentin. High-plasticity organoids showed increased invasive outgrowth and Vimentin expression. Scale bars, 100 μm. **E.** Representative immunofluorescence images and quantification of tumor organoids treated with vehicle control or trametinib. MEK inhibition reduced invasive outgrowth and Vimentin expression.

We next asked when EMT-associated plasticity is acquired during tumor progression. Independent tumors arising within the same mouse differed in E-lo abundance, indicating that systemic host factors alone do not determine EMT activation in C3-TAg tumors **(Fig. S3C)**. We therefore examined how tumor cell state influenced the capacity to undergo EMT-associated phenotypic transitions. Hybrid EMT states, such as those detected along the EMT trajectories in >15 mm tumors, have been proposed to confer plasticity that permits further progression toward mesenchymal-like states. However, E-hi cells isolated from 5 mm tumors, which lacked these canonical hybrid states, also readily generated E-lo cells in organoid culture. Thus, tumor cells can acquire the capacity to transition to a mesenchymal-like state before overt hybrid states emerge **(Fig. 3C)**. In contrast, organoids derived from hyperplastic precursor lesions, which express the C3-TAg oncogene, showed the lowest frequency of E-lo emergence **(Fig. 3C)**.

The greater generation of E-lo cells by 5 mm tumor organoids was accompanied by increase invasion and Vimentin expression compared with hyperplasia-derived organoids **(Fig. 3D)**. MEK1/2 inhibition with trametinib reduced both Vimentin induction and invasion **(Fig. 3E)**. Together, these findings suggest that progression-associated differences in EMT plasticity reflect, at least in part, changes in cell-intrinsic response to ERK1/2 pathway activation. Importantly, the low EMT-associated plasticity of hyperplasia-derived organoids did not reflect poor tumor-initiating capacity. These organoids formed orthotopic tumors with kinetics comparable to those of organoids derived from more advanced lesions **(Fig. S3D)**. Thus, functional EMT-associated plasticity emerges as a distinct property during tumor progression, before overt hybrid EMT states become established, and is separable from tumor-initiating potential.

Together, these findings show that EMT progression in C3-TAg tumors is directional, heritable, and acquired during tumor evolution. Early oncogenic transformation produces epithelial-like cells with limited EMT plasticity, whereas later tumor stages give rise to hybrid E-hi cells capable of transitioning toward stable mesenchymal-like E-lo endpoints. These E-lo populations as comparatively stable mesenchymal-like states that persists across in vitro and in vivo contexts, providing a functional mechanism for how EMT plasticity generates and maintains phenotypic heterogeneity in advanced tumors.

### Metastatic competence is shared across hybrid and mesenchymal-like EMT states

We next asked how EMT heterogeneity influences metastatic behavior. Prior studies in breast, pancreatic, and squamous carcinomas have suggested that E-lo mesenchymal-like cells represent the principal metastatic population, particularly in tumors with robust EMT induction (5, 27, 59, 60). We therefore tested, in our physiologically heterogeneous system, whether hybrid E-hi and mesenchymal-like E-lo states differ in their ability to seed and grow metastatic lesions.

To compare metastatic competence across EMT states, we first used serial transplantation to generate multiple independently derived C3-TAg tumor lines that naturally evolved enrichment for either E-hi or E-lo cells during passaging **(Fig. 4A, S4A)**. Tail-vein experimental metastasis assays showed that both E-hi- and E-lo-enriched tumor lines were robustly metastatic, indicating that metastatic competence was not restricted to the mesenchymal-like E-lo state **(Fig. 4B–D)**. To test whether E-hi cells could colonize organs beyond the lung, we performed intracardiac injection. E-hi cells formed liver metastases in addition to lung metastases; lung metastases were detected in all injected mice, whereas liver metastases were detected in 3 of 5 mice **(Fig. S4B–C)**. In some animals, liver metastatic burden exceeded lung metastatic burden within the same mouse. Thus, E-hi cells can colonize multiple distant organs, indicating that their metastatic potential is not restricted to pulmonary colonization **(Fig. S4B–C)**. We next directly compared FACS-isolated E-hi and E-lo populations from primary C3-TAg tumors in tail-vein metastasis assays across three independent models (Fig. 4E). E-hi cells formed metastases in all three models **(Fig. 4E–G, S4D)**. In two models, E-hi cells produced greater metastatic burden than E-lo cells, whereas in the third model E-hi cells were modestly less metastatic than their E-lo counterparts **(Fig. 4E–G, S4D)**. These findings indicate that mesenchymal-like E-lo cells are not uniquely enriched for metastatic competence. Rather, metastatic potential can be distributed across EMT states, including both hybrid E-hi and mesenchymal-like E-lo cells.

**Figure 4.**
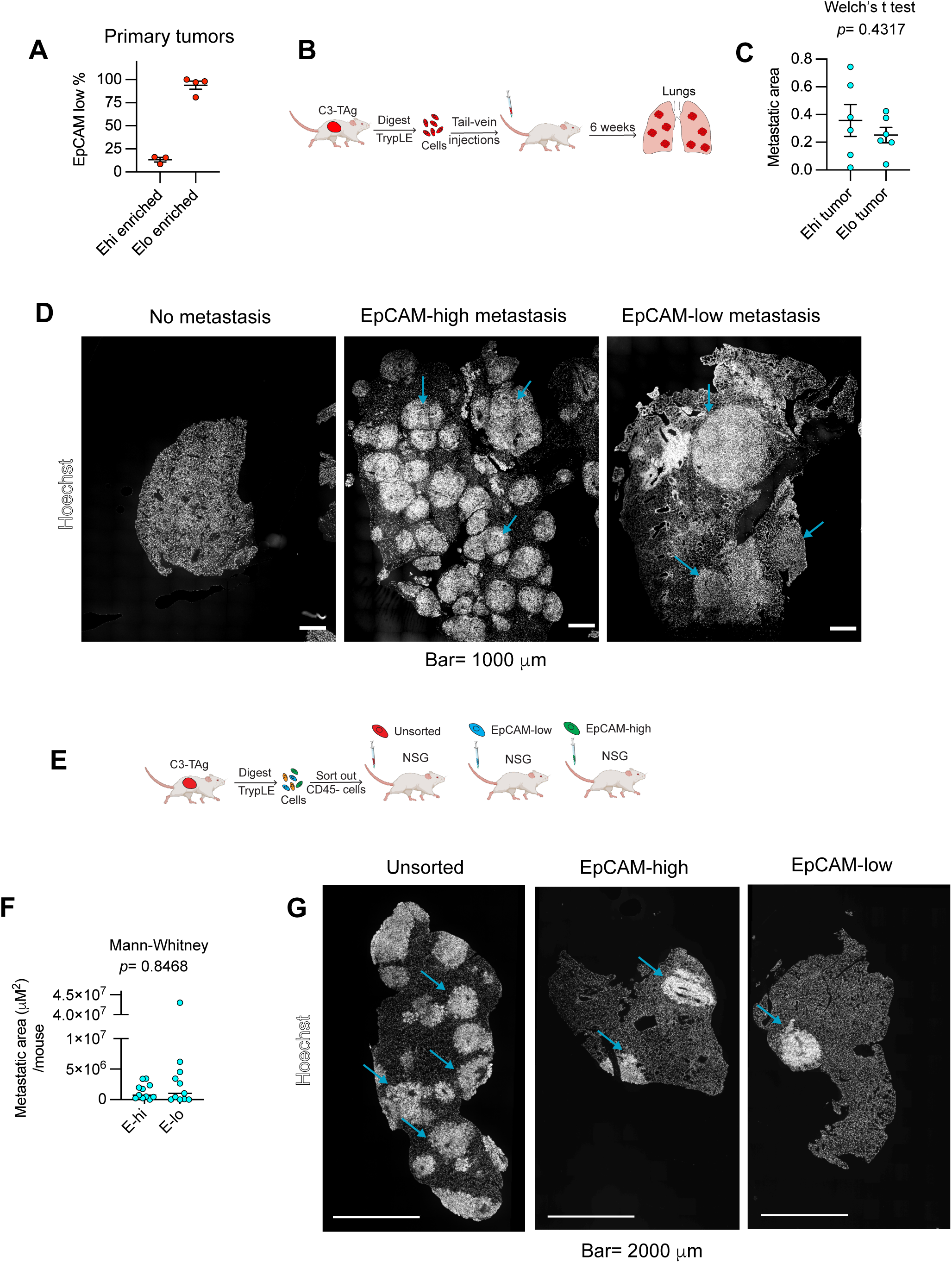
Hybrid EpCAM-high and mesenchymal-like EpCAM-low C3-TAg tumor cells are both metastatically competent. **A.** FACS analysis showing the differential EpCAM-low cell abundance in the injected tumor lines. **B.** Experimental workflow for tail-vein injection of independently derived C3-TAg tumor lines enriched for EpCAM-high or EpCAM-low states. **C.** Quantification of metastatic area per mouse. **D.** Representative Hoechst-stained lung sections show metastatic burden from EpCAM-high-and EpCAM-low-enriched tumor lines. Arrows indicate metastatic lesions. Scale bars, 1000 μm. **E.** Experimental workflow for direct FACS isolation of CD45-negative unsorted, EpCAM-high, and EpCAM-low C3-TAg tumor cells followed by transplantation into NSG mice for lung metastasis assays. **F.** Quantification showing that both EpCAM-high and EpCAM-low tumor cell populations formed lung metastases. **G.** Representative Hoechst-stained lung sections and. Arrows indicate metastatic lesions. Scale bars, 2000 μm.

### EMT state influences the phenotype of metastatic lesions

We next asked whether the EMT state of the metastasis-initiating population influences the phenotype of the resulting metastatic lesions. If mesenchymal-to-epithelial transition (MET) were obligate during metastatic colonization (13, 16, 60, 61), metastases initiated by hybrid E-hi and mesenchymal-like E-lo populations would be expected to converge toward a similar epithelial or hybrid phenotype.

To test this prediction, we analyzed Vimentin expression and cell morphology in metastatic lesions derived from E-hi- and E-lo-enriched tumor lines **(Fig. 5A)**. Metastases derived from E-lo-enriched tumor lines showed widespread Vimentin expression, spindle-like morphology, and invasive growth within the lung **(Fig. 5B–C)**. In contrast, metastases derived from E-hi-enriched tumor lines showed little or sporadic Vimentin expression and retained epithelial morphology with glandular architecture **(Fig. 5B–C)**. Similar differences were observed in metastases initiated by FACS-isolated E-hi and E-lo cells, indicating that the phenotype of the initiating population influences metastatic phenotype **(Fig. S5A–C)**.

**Figure 5.**
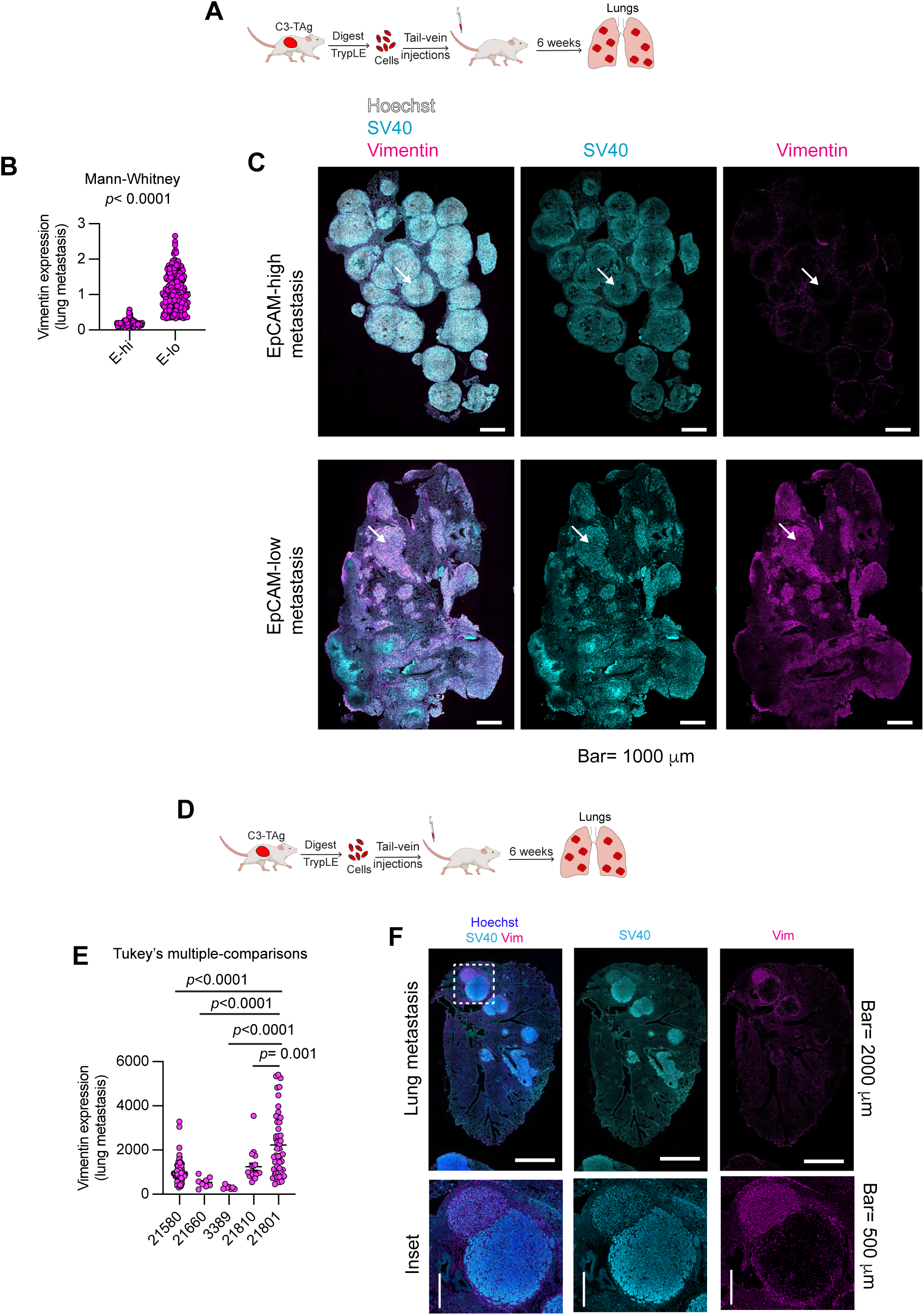
Metastatic lesions derived from EpCAM-high and EpCAM-low C3-TAg tumor cells retain distinct EMT-associated phenotypes. **A.** Experimental workflow for tail-vein injection of independently derived C3-TAg tumor lines followed by analysis of lung metastases. **B.** Quantification of Vimentin expression in lung metastases generated by EpCAM-high- and EpCAM-low-enriched tumor lines. **C.** Representative immunofluorescence images of lung metastases derived from EpCAM-high-or EpCAM-low-enriched tumor lines stained for Hoechst, SV40, and Vimentin. Arrows indicate Vimentin-positive regions. Scale bars, 1000 μm. **D.** Experimental workflow for tail-vein injection of independently derived C3-TAg tumor lines followed by analysis of lung metastases. **E.** Quantification of Vimentin expression across lung metastases generated from independent spontaneous C3-TAg tumors. **F.** Representative immunofluorescence images showing heterogeneous Vimentin expression within lung metastases generated from C3-TAg tumor lines. Images show Hoechst, SV40, and Vimentin staining with enlarged insets. Scale bars, 2000 μm for whole-lung images and 500 μm for inset images.

We also examined metastases arising after tail-vein injection of new spontaneous C3-TAg primary tumors **(Fig. 5D)**. Many metastases displayed epithelial morphology, glandular architecture, and low Vimentin expression, whereas others were mesenchymal-like with high Vimentin expression **(Fig. 5E–F, S5D–E)**. Epithelial-like and mesenchymal-like metastases could be found in close proximity within the same lung, arguing that the local metastatic microenvironment alone does not determine metastatic EMT phenotype **(Fig. 5E–F, S5D–E)**. These findings are consistent with the EMT phenotype of metastases being influenced by the intrinsic state of the metastasis-initiating population and indicate that a single primary tumor can generate pheno-typically distinct classes of metastases.

Collectively, these findings reveal that metastatic lesions do not converge uniformly on a canonical epithelial state. Although some plasticity may occur during metastatic outgrowth, hybrid E-hi and mesenchymal-like E-lo populations produce distinct metastatic phenotypes. Thus, EMT state does not dictate metastatic competence—both hybrid and mesenchymal-like tumor cells can seed metastases—but it strongly influences the phenotype of the metastatic lesions that arise. Together, these results support a model in which EMT-associated heterogeneity is generated by acquisition of plasticity and maintained through stable mesenchymal-like commitment, while metastatic competence remains distributed across EMT states that produce distinct classes of metastatic lesions.

## DISCUSSION

Our findings define a temporal and functional framework for how EMT-associated heterogeneity emerges during TNBC progression. Tumor cells first acquire increased EMT-associated plasticity before overt hybrid or mesenchymal-like states become abundant. This increased in intrinsic plasticity is followed by diversification through at least two transcriptionally distinct EMT programs that share early loss of mammary lineage and epithelial features but differ in downstream regulatory composition and mesenchymal gene expression. Finally, both hybrid and mesenchymal-like states retain metastatic competence, while the state of the initiating population influences the phenotype of the resulting metastatic lesions. These findings distinguish the acquisition of EMT plasticity, progression toward stable mesenchymal-like states, and metastatic competence as related but separable features of tumor evolution (6, 61).

A central result is that increased EMT-associated plasticity emerges before overt hybrid EMT states are readily detectable. Cells isolated from 5-mm tumors readily generated E-lo progeny in organoid culture, despite the limited representation of overt hybrid and mesenchymal-like states at this stage. In contrast, organoids derived from hyperplastic precursor lesions showed substantially less E-lo emergence while retaining robust tumor-initiating capacity. Prior studies have shown that EMT responsiveness can be influenced by the cell of origin or enhanced experimentally through alteration of chromatin and histone-modification states (60, 62, 63). Our findings extend this work by showing that increased EMT responsiveness is acquired at a defined stage of spontaneous tumor progression, between oncogene-expressing hyperplasia and early invasive disease.

Pseudotime analysis identified early loss of mammary lineage-associated genes and cell–cell junction components before broad induction of mesenchymal genes. These findings suggest that weakening of lineage identity and epithelial organization may create a permissive state in which tumor cells respond more readily to EMT-inducing signals. Loss or relaxation of lineage restriction has similarly been linked to increased phenotypic plasticity during tissue repair and tumor progression (64–67). Although the functional organoid studies and transcriptional trajectory analysis do not directly assign increased plasticity to a specific pseudotime position, these complementary results support a model in which functional competence for EMT precedes overt epithelial–mesenchymal marker co-expression. Determining whether loss of individual mammary lineage regulators directly promotes this permissive state will require targeted perturbation and longitudinal analysis.

The asymmetric transition behavior of E-hi and E-lo populations further explains how EMT heterogeneity can be generated and maintained. E-hi cells consistently produced E-lo progeny, whereas E-lo cells rarely regenerated an E-hi population. Moreover, tumors initiated from E-lo organoids retained a predominantly E-lo phenotype in vivo. EMT-associated states therefore differ substantially in reversibility, consistent with evidence that hysteresis and cellular memory can stabilize particular EMT-associated phenotypes after withdrawal of an inducing signal in experimental model systems (16, 18, 25). The expression profile of E-lo cells was also consistent with components of previously described EMT-stabilizing regulatory circuits, including increased ZEB-family transcription factor expression, reduced epithelial character, and induction of mesenchymal genes (16). Thus, hybrid-like E-hi populations retain considerable transition capacity, whereas E-lo populations represent comparatively stable mesenchymal-like states. This asymmetric organization provides a mechanism through which plastic tumor populations can progressively generate persistent phenotypic heterogeneity (68, 69).

Our single-cell analysis also shows that EMT-associated diversification cannot be fully represented by a single epithelial-to-mesenchymal axis. The inferred trajectories shared early reductions in mammary lineage genes, epithelial junctional components, EpCAM, and E-Cadherin, but diverged in EMT transcription factor usage and mesenchymal-associated effector programs. Other tumor models have similarly revealed context-dependent or parallel EMT programs rather than a single conserved transcriptional sequence (11, 59, 70). The p63-positive Cluster 8 population was positioned separately from the principal bifurcating structure, suggestting that not all hybrid states are obligatory intermediates leading toward a common mesenchymal endpoint. Instead, advanced C3-TAg tumors contain multiple hybrid and mesenchymal-like states with distinct transcriptional organizations, consistent with the broader diversity of EMT transition states identified across tumor types (7).

Our results suggest that differential engagement of ERK1/2-regulated transcription distinguishes the two EMT trajectories. The ERK1/2-low trajectory progressed through hybrid states in which Twist1 and Vimentin expression began to increase, followed by further loss of epithelial character and induction of Zeb1, Prrx1, and a distinct cohort of extracellular matrix reorganization genes. Both trajectories showed enrichment for TGFβ-induced programs, particularly those associated with prolonged TGFβ exposure, indicating that TGFβ-responsive transcription contributes to each branch. In contrast, the ERK1/2-high trajectory showed additional enrichment for ERK1/2-regulated genes, including extracellular matrix-interaction genes and the autocrine EGFR ligands Areg and Ereg. Although both trajectories showed progressive loss of epithelial features and induction of shared mesenchymal genes, including Vimentin and Zeb2, they differed in EMT transcription factor usage and extracellular matrix regulatory programs. This distinction is consistent with our previous finding that EGF-driven ERK1/2 activation modifies the transcriptional response to TGFβ and restricts the extent of TGFβ-induced mesenchymal and extracellular matrix gene expression (22). Together, these findings suggest that TGFβ- and ERK1/2-regulated signals do not define completely separate EMT pathways but instead are differentially integrated to generate transcriptionally distinct EMT-associated states. This interpretation is consistent with the context-dependent and combinatorial regulation of EMT by TGFβ, RAS–ERK, and receptor tyrosine kinase signaling pathways (11, 12, 71, 72).

This organization of EMT-associated heterogeneity into multiple trajectories also has implications for how EMT states are defined. Vimentin induction and E-Cadherin loss occurred relatively late along both trajectories, after disruption of mammary lineage and junctional programs. Cluster 2 retained substantial epithelial character despite expressing an EMT-enriched program and genes associated with matrix interaction and invasion. Standard classification based on Vimentin, E-Cadherin, or a limited set of canonical EMT transcription factors may therefore fail to detect early plastic states or distinguish functionally different hybrid populations (6, 19, 73). More complete characterization will require integration of lineage identity, signaling context, transcriptional state, and functional transition capacity (7, 13).

Our metastasis experiments further separate EMT phenotype from metastatic competence. Both E-hi- and E-lo-enriched tumor lines formed robust metastases, and directly isolated E-hi populations generated metastases in all three models tested. Although the relative metastatic burden produced by E-hi and E-lo cells varied among tumors, neither population was consistently superior. This result differs from studies in which metastatic activity was concentrated within hybrid or more mesenchymal EMT-associated populations (5, 26, 59, 60, 74). These differences may reflect the composition of EMT states represented in each model. For example, MMTV-PyMT tumors predominantly disseminate through Krt14-positive hybrid-EMT clusters and contain relatively few fully mesenchymal cells (22, 26, 38, 75, 76). Conversely, studies of models dominated by E-lo or mesenchymal-like populations have often identified the greatest metastatic activity within subsets that retain marginally greater epithelial character within the broader E-lo population (5, 59). In contrast, C3-TAg tumors contain abundant and transcriptionally diverse hybrid and mesenchymal-like populations (22, 38). By examining tumors spanning this broader EMT spectrum, our study shows that metastatic competence is not restricted to either the most mesenchymal-like population or a single preferred intermediate state. Instead, it is distributed across multiple EMT-associated populations, with the relative metastatic fitness of E-hi and E-lo cells varying among tumors. This conclusion is consistent with studies showing that distinct EMT-associated states can each contribute to metastatic progression in a context-dependent manner (26, 27, 38, 59). The ability of E-hi cells to colonize both lung and liver further indicates that their metastatic potential is not confined to a single organ environment.

Although EMT state did not uniquely determine metastatic competence, it strongly influenced the phenotype of the lesions that formed. Metastases derived from E-lo-enriched or FACS-iso^-^ lated E-lo populations generally retained widespread Vimentin expression, spindle-like morphology, and invasive growth. In contrast, E-hi-derived metastases more often displayed epithelial morphology, glandular organization, and little or sporadic Vimentin expression. Phenotypically distinct metastases also occurred near one another within the same lung following injection of cells from spontaneous primary tumors, arguing that the local metastatic environment alone does not impose a uniform EMT phenotype. Instead, metastatic colonization is likely shaped by interactions between the intrinsic state of disseminated tumor cells and the signals encountered in the metastatic niche (40, 77).

The persistence of Vimentin-high, mesenchymal-like lesions demonstrates that metastatic outgrowth does not universally require complete mesenchymal-to-epithelial transition. Previous lineage-tracing and genetic studies have questioned whether EMT is obligatory for metastatic dissemination and have suggested that epithelial or partially transitioned cells can efficiently form metastases (26, 78–80). Our findings extend this framework by showing that mesenchymal-like E-lo populations can also produce macrometastatic lesions while retaining prominent mesenchymal-associated features. Conversely, the epithelial organization of many E-hi-derived metastases does not necessarily indicate that these cells underwent MET, because they may have disseminated without first adopting a strongly mesenchymal phenotype. The phenotype of a metastatic lesion therefore reflects both the state of the initiating population and any subsequent plasticity during colonization. These findings refine models in which MET promotes metastatic colonization and provide a potential explanation for why many clinical metastases appear epithelial despite extensive evidence that EMT-associated programs promote dissemination (77, 81).

The persistence of distinct metastatic phenotypes may also have therapeutic consequences. Epithelial-like and mesenchymal-like metastases could differ in signaling dependencies, stromal interactions, and treatment response. EMT-associated plasticity and state heterogeneity have been linked to therapeutic adaptation, immune regulation, and resistance across multiple cancer settings (4, 7, 82, 83). Therapies directed at individual EMT-inducing pathways may therefore suppress only subsets of metastasis-competent cells. Identifying vulnerabilities shared across EMT states, as well as dependencies specific to stable mesenchymal-like populations, will be important for developing strategies that remain effective across heterogeneous metastatic disease.

Together, our findings support a model in which increased EMT-associated plasticity precedes overt state diversification, distinct regulatory programs generate multiple hybrid and mesenchymal-like populations, and metastatic competence remains distributed across this phenotypic landscape. EMT-associated heterogeneity therefore produces multiple cellular states capable of metastatic colonization and distinct metastatic outcomes rather than a single optimal metastasis-initiating state or obligatory epithelial endpoint.

## MATERIALS AND METHODS

**Mice**. The C3-TAg [FVB-Tg(C3-1-TAg)cJeg/Jeg], PyMT [FVB/N-Tg(MMTV-PyVT)634Mul/J] and NOD.Cg-Prkdcscid Il2rgtm1Wjl/SzJ (NSG) mice were purchased from The Jackson Laboratory (C3-TAg: 013591, PyMT: 002374 and NSG: 005557) and (**Supplementary file 1**). Mice were housed, bred and euthanized in accordance with a protocol approved by the Institutional Animal Use and Care Committee at Georgetown University (IACUC protocol 2017-0076) and in compliance with the NIH Guide for the Care and Use of Laboratory Animals. Female mice were used for all experiments.

### Tumor organoid derivation

The largest tumors from female C3-TAg and PyMT mice were minced and dissociated for up to 120 min at 37°C in a mixture of 1 mg/ml Collagenase (Worthington, LS004188), 20 U/ml DNase (Invitrogen, 18047-019), 5% Fetal Bovine Serum (FBS) (Peak Serum, PS-FB2), ITS (Lonza, 17-838Z), non-essential amino acids (Sigma, M7145) and 10 ng/ml FGF-2 (Peprotech, 100-18C) in DMEM/F-12 (Corning, 10-092-CV). Dissociated tumors were pelleted at 80 x *g* for 2 min and the supernatant was discarded. Tumor organoids were then rinsed up to 5 times in 5% FBS in DMEM/F-12 followed by filtering through a 250 µm tissue strainer. Organoids were then tested in invasion assays or to establish organoid lines and cell lines (**Supplementary file 1**).

### Cell and organoid culture

Cell and organoid lines were generated from C3-TAg tumors **(Supplementary file 1)** and tested for mycoplasma (Lonza, LT07-703) prior to use and the creation of frozen stocks. Cells and organoids were routinely used within 25 passages. Culture conditions for all cell and organoid lines are summarized in **Supplementary file 1.** Organoids were passaged at least once per week by dissociating tumor organoids into single cell suspensions with Dispase (Sigma, SCM133) and TryPLE (Gibco, 12605-010). The cell suspensions were plated at a density of 200,000-500,000 cells per well in a 24-well ultra-low adhesion plate (Sigma, CLS3474) to form multicellular aggregates for at least 16 h. The aggregates were then embedded in Matrigel (Corning, 354230) or Cultrex (Biotechne, 3533-005-02) and overlaid with Organoid Media (**Supplementary file 1**) in 6-well ultra-low adhesion plates (Sigma, CLS3471).

### Orthotopic Tumor Models

For orthotopic transplantation experiments, organoid cells were first cultured in Organoid media. To initiate orthotopic tumors, 250,000 organoid cells were injected in the right 4^th^ fat pad of 8–12 week-old female NSG mice. Mice were euthanized when tumors reached a maximum allowed diameter of 2 cm, and tissue collection was performed. Representative portions of primary tumors were fixed in formalin followed by paraffin embedding and sectioning by Histology and Tissue Shared Resource (HTSR) at Georgetown University.

### Organoid invasion

Organoids in 30 µl of ECM were plated onto a 20 µl base layer of ECM in 8-well chamber slides (Falcon, 354108) and allowed to invade for 48 h unless otherwise indicated. The ECM was a mixture of 2.4 mg/ml rat tail collagen I (Corning, CB-40236) and 2 mg/ml of reconstituted basement membrane (Cultrex) unless otherwise indicated. Tumor organoids were analyzed in a base medium of DMEM/F-12, ITS and FGF. Organoids were fixed and stained with Hoechst and Phalloidin. Images were acquired with a Zeiss LSM800 laser scanning confocal microscope using 10x/0.45 (Zeiss, 1 420640-9900-000) or 20x/0.8 (Zeiss, 1 420650-9902-000) objectives. To determine the area of invasion, an image mask for each organoid was generated using ImageJ (NIH) based on the Hoechst or Phalloidin signal. The total area of the invading cell nuclei was determined using the “Measure” function in ImageJ. Circularity was quantified using the “Analyze Particle” function.

### Immunofluorescence and immunoblotting

Experiments and analysis were performed as described (21) using antibodies detailed in **Supplementary file 1.** Immunoblots were imaged using an Odyssey scanner (Licor, 9120). Immunofluorescence images were acquired using 10x/0.45 (Zeiss, 1 420640-9900-000) and 20x/0.8 (Zeiss, 1 420650-9902-000) objectives. Images were exported as TIFFs and analyzed with ImageJ. The freehand selection tool was used to define ROIs containing individual organoids for analysis. Vimentin and E-Cadherin expression were analyzed in 5 µm histology sections. Organoids in 8-well chamber slides (Falcon, 354108) were washed 3x with distilled water. After detaching the chamber walls from the slides, the organoid/ECM gels were gently picked up using a single edge blade (Personna, 94-120-71) and laid out on a strip of parafilm (Bemis, PM-999). The 8-well chamber walls were placed around the gels and 150-200 µl of hydroxyethyl agarose processing gel (Thermo Fisher, HG 4000-012) was poured into the well. After the hydroxyethyl agarose processing gel solidified, it was transferred into histosettes (Simport, M498-3) and stored in 70% EtOH until embedding in paraffin by the Georgetown Histology and Tissue Shared Resource. The 5 µm sections were deparaffinized and immunostained as described (15) to detect Vimentin and E-Cadherin expression (**Supplementary file 1)**. For individual organoids, Vimentin expression was defined as the “Mean Gray Value” using the “Measure” function in ImageJ using ImageJ.

### Immunohistochemistry

Sample processing and staining was performed as described (84). Images were acquired using 10x/0.45 (Zeiss, 1 420640-9900-000) and 20x/0.8 (Zeiss, 1 420650-9902-000) objectives. The area of metastasis was determined by dividing the area of metastases in each lung section by the total area of lung tissue in the section. Zen Blue software was used to define image masks based on SV40 or Vimentin signal. The Vimentin-positive area was divided by the total SV40-positive area to quantify relative Vimentin expression in metastatic lesions.

### Flow cytometry

Cells were dissociated with TrypLE (Gibco, 12605-010) into single cell suspensions, and then washed 2x with FACS buffer (PBS + 2% FBS). Cells were then incubated with anti-CD326 (EpCAM)-PE/Dazzle 594 (Biolegend, 118236) at 4°C for 30 minutes. After staining, cells were washed with FACs buffer twice, and stained with Helix NP Blue (Biolegend, 425305) in order to exclude dead cells. Cells were then subjected to flow cytometry analysis using a BD LSRFortessa flow cytometer. Results were analyzed using FCSExpress 7 (De Novo Software).

### Single cell gene expression analysis

Libraries were prepared with the Georgetown Genomics and Epigenomics Shared Resource using the Chromium Next GEM Single Cell 3’ kit following the manufacturer’s protocol (10X Genomics). Targeted cell recovery was 5,000 cells per sample. Libraries were sequenced by Novogene with the HiSeq PE150 platform at a depth of approximately 70 million paired-end 150 base pair reads per sample. Cell Ranger software was used for demultiplexing samples, barcode processing and alignment to the mouse reference genome (mm10). Cells were subjected to quality control and filtering based on per-cell total transcript counts and mitochondrial transcript abundance (85). Uniform Manifold Approximation and Projection (UMAP) dimensionality reduction and clustering were performed in R using Seurat (86). Clusters of non-tumor cells were determined based on gene expression using canonical lineage markers. Pseudotime trajectory analysis and gene module derivation was performed with Monocle3 (58).

### Statistical Methods

Data were analyzed with Prism 9.0.2 (GraphPad). Data with a normal distribution, as determined by Shapiro-Wilk test, was analyzed by two-tailed Student’s t-test. Data that did not pass a normality test were analyzed by Mann-Whitney U test or Kruskal-Wallis test with Dunn’s Multiple-comparison test. The specific statistical tests are indicated in the figure legends. Sample numbers are defined and indicated in the figure legends or on the figure panels.

## AUTHORS’ CONTRIBUTIONS

**F. Alsharief:** Conceptualization, investigation, writing-review and editing. **R.K. Suter:** Conceptualization, investigation writing-review and editing. **A. Nasir:** Conceptualization, investigation and editing. **I. Cruz:** investigation. **G.W. Pearson:** Conceptualization, investigation, writing-review and editing, funding acquisition.

## ACKNOWLEDGEMENTS

Work was supported by NIH R01CA218670, Georgetown Women and Wine (to G.W. Pearson), and NIH P30CA051008.

---

**Figure S1.**
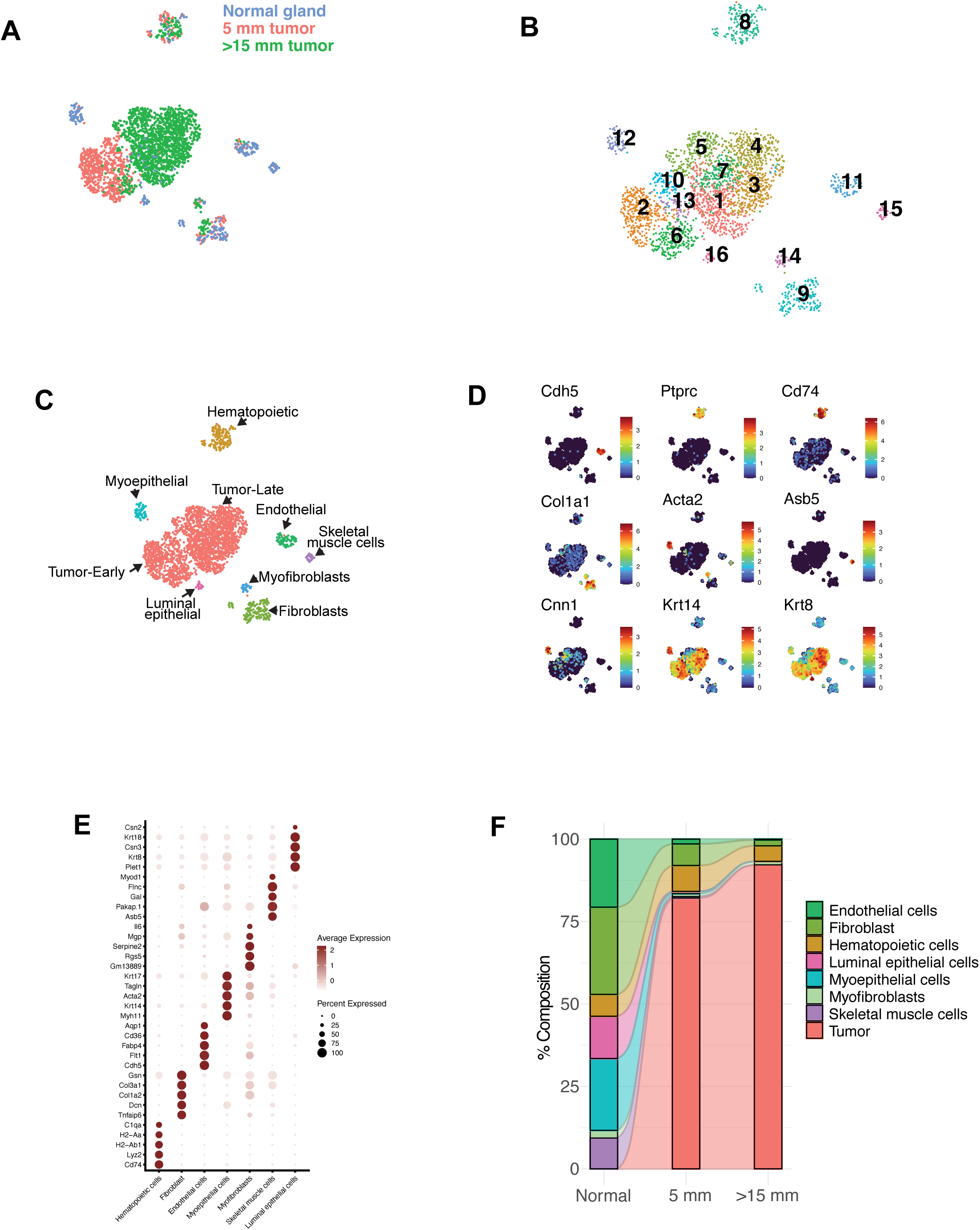
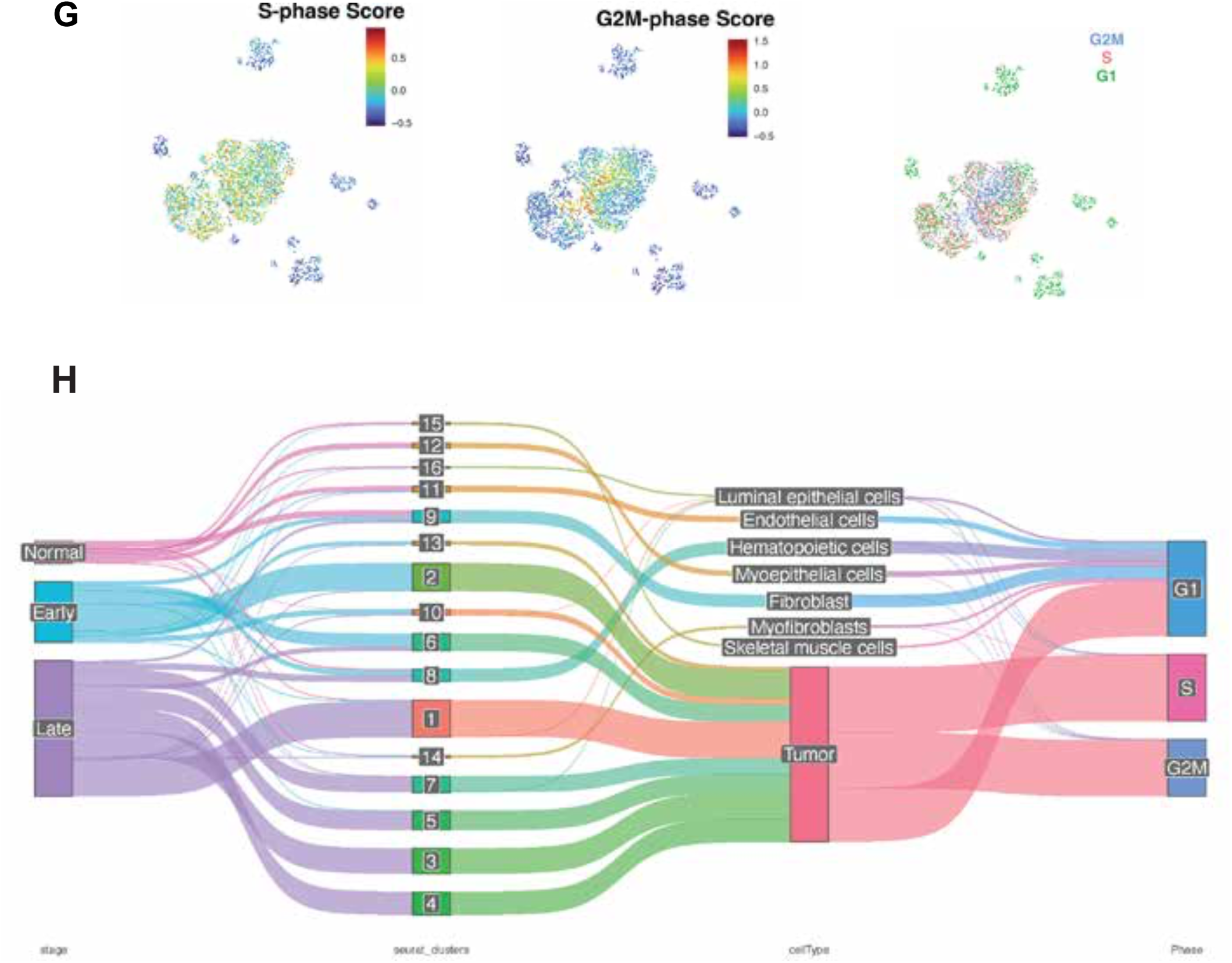
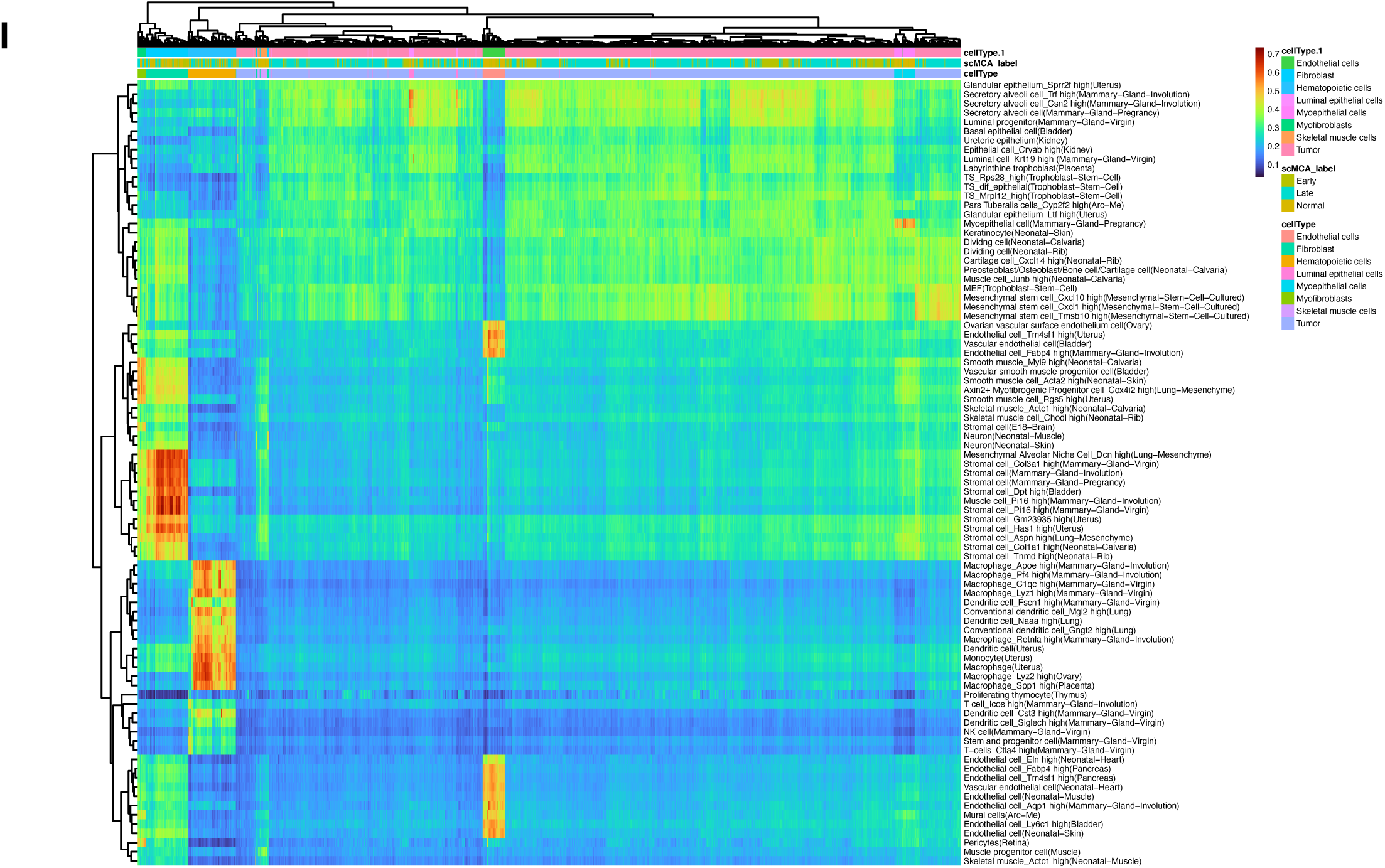
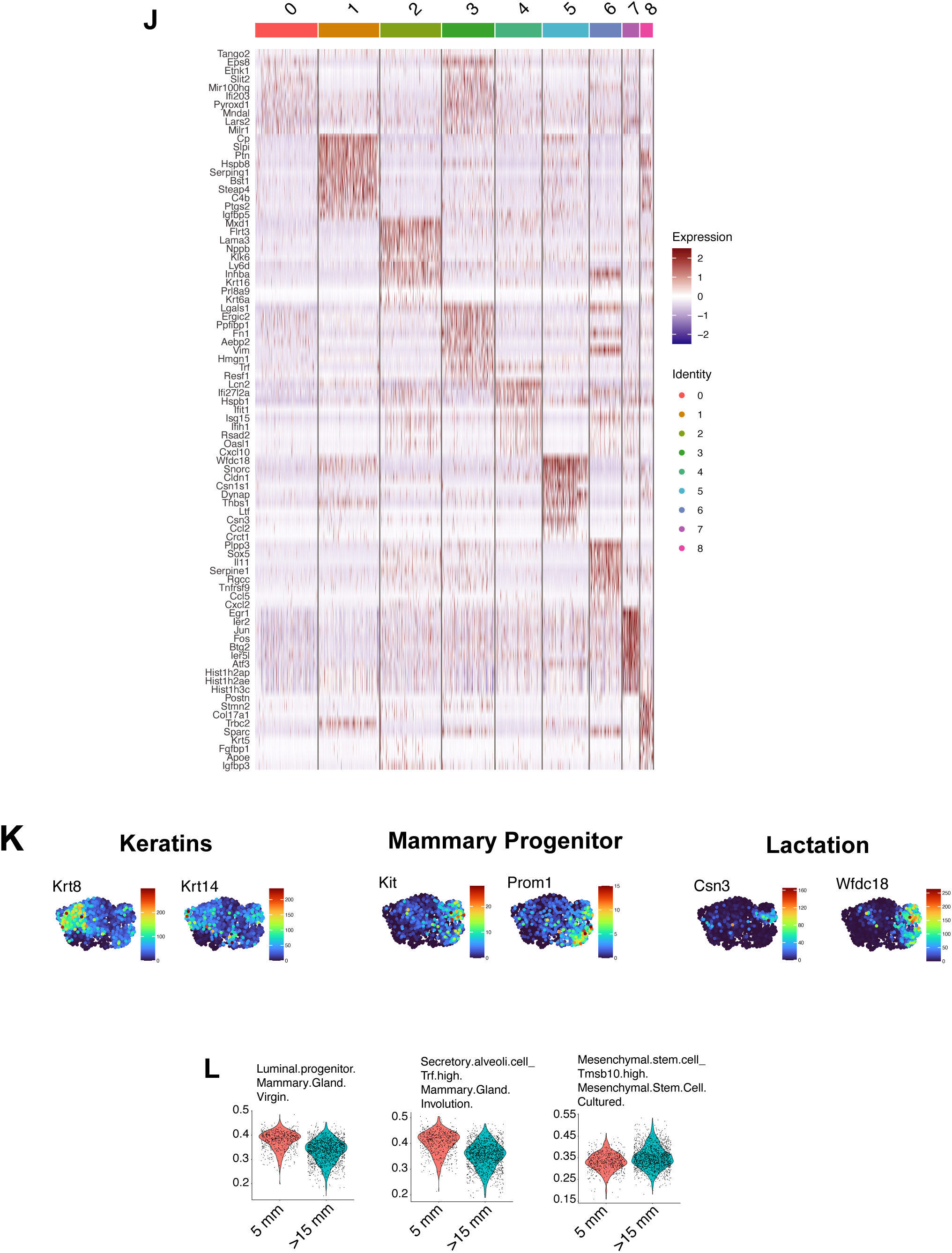
Related to C3-TAg tumor progression generates extensive EMT-associated transcriptional heterogeneity. **A.** UMAP showing cells from normal mammary gland, 5 mm and >15 mm tumors annotated by sample source. **B.** UMAP showing cell clusters from normal mammary gland, 5 mm and >15 mm tumors annotated by Seurat cluster. **C.** UMAP showing cells from normal mammary gland, 5 mm and >15 mm tumors annotated by cell type. **D.** UMAP showing the expression of cell lineage specific genes in cells from normal mammary gland, 5 mm and >15 mm tumors annotated by cell type. **E.** Bubble plot showing the expression of marker genes in the indicated normal cell lineages. **F.** Alluvial stacked bar plot showing the percentage of indicated cell types in normal mammary gland, 5 mm and > 15 mm tumors. **G.** UMAPs showing the cell cycle phase of cells from normal mammary gland, 5 mm and >15 mm tumors annotated by sample source. **H.** Alluvial plot showing the distribution of cells from normal mammary gland, 5 mm and >15 mm tumors. **I.** Heatmap showing the similarity of the indicated cells to normal cell types from a mouse scR-NA-seq atlas. **J.** Heatmap showing the expression of top genes differentially expressed between clusters. **K.** UMAPs showing the expression of representative luminal progenitor, lactation-associated and basal and genes in 5 mm and >15 mm tumor cells. **L.** Quantification compates the similarity of 5 mm and >15 mm tumor cells to luminal progenitor, secretory alveolar and mesenchymal stem cells.

**Figure S2.**
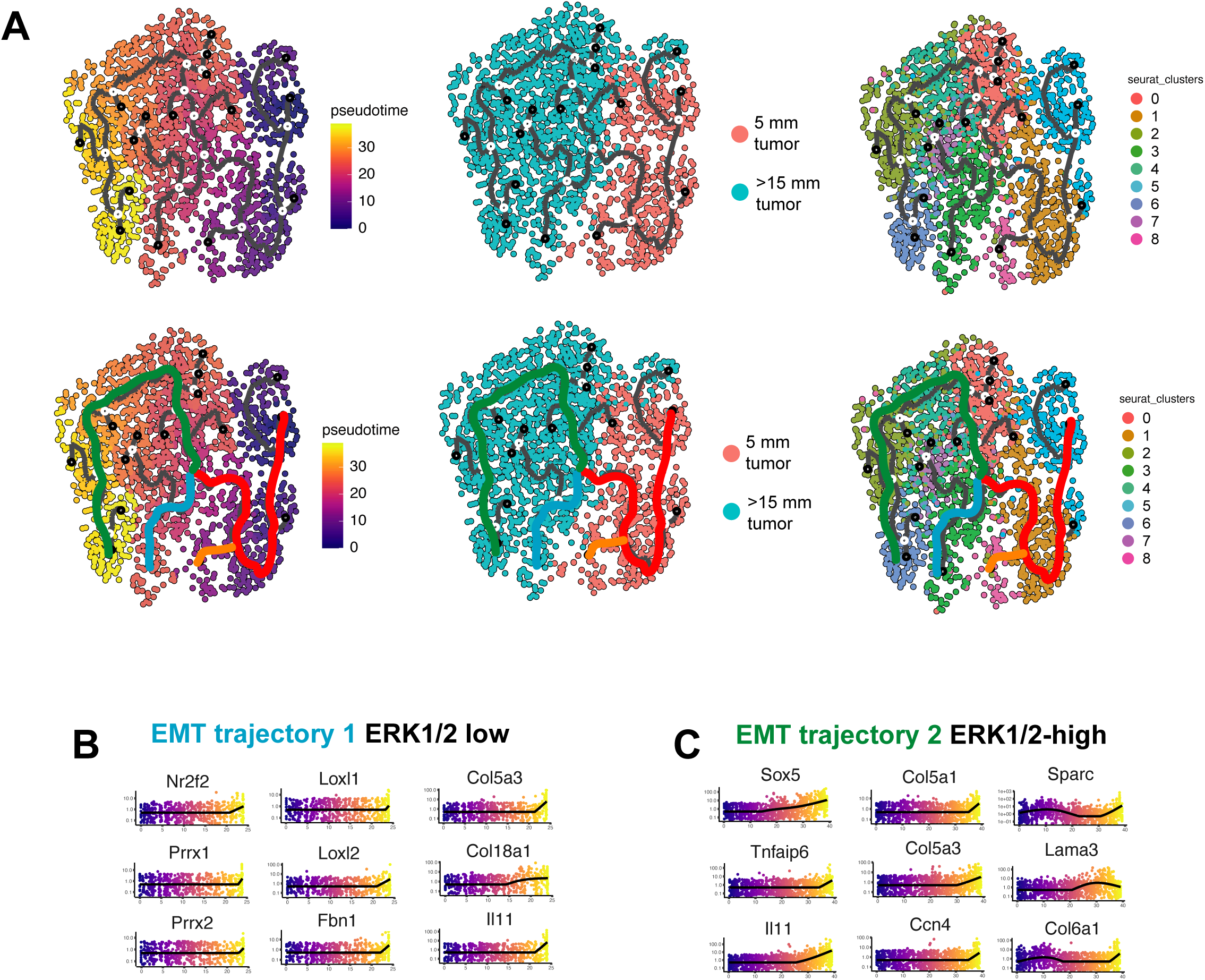
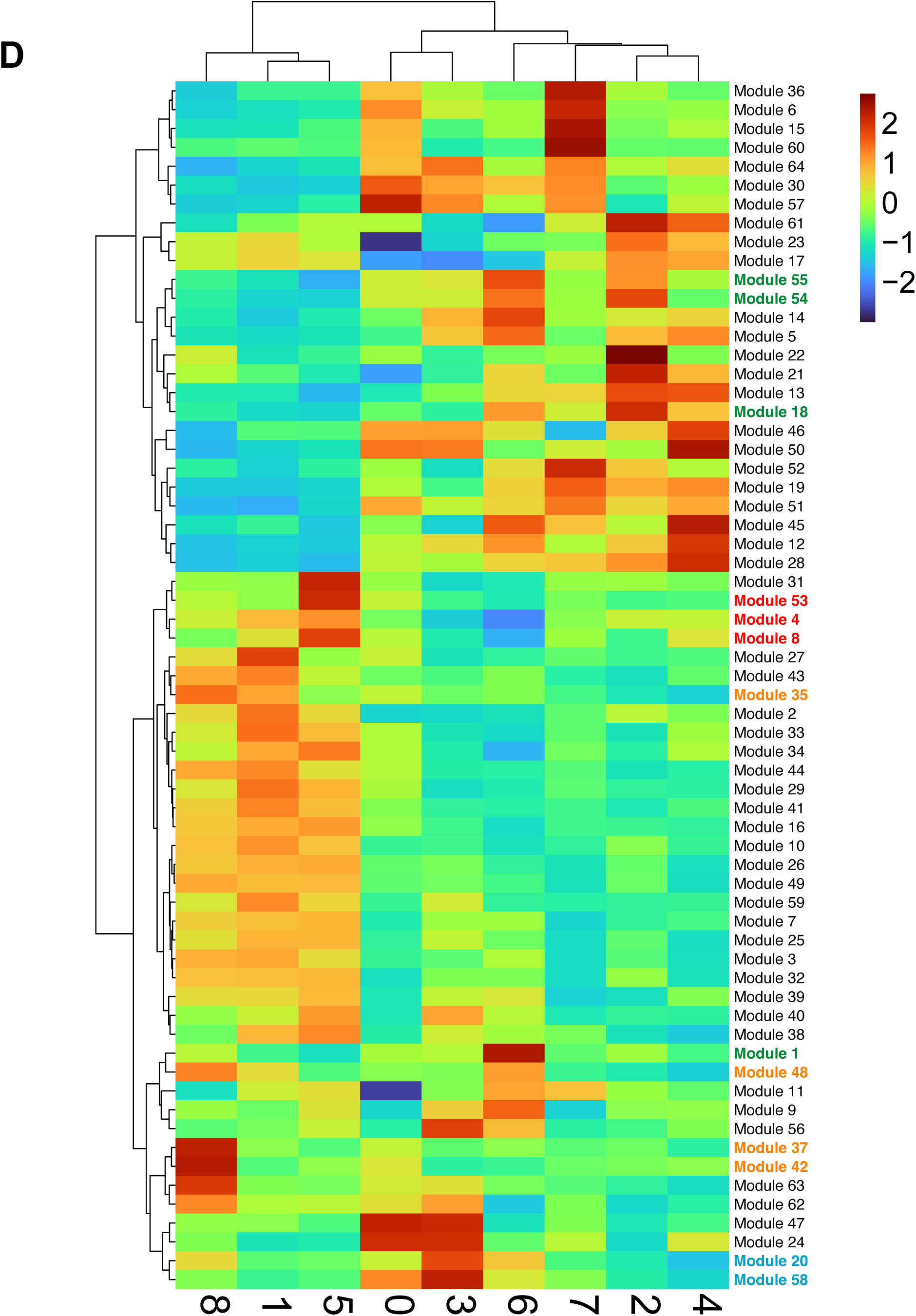
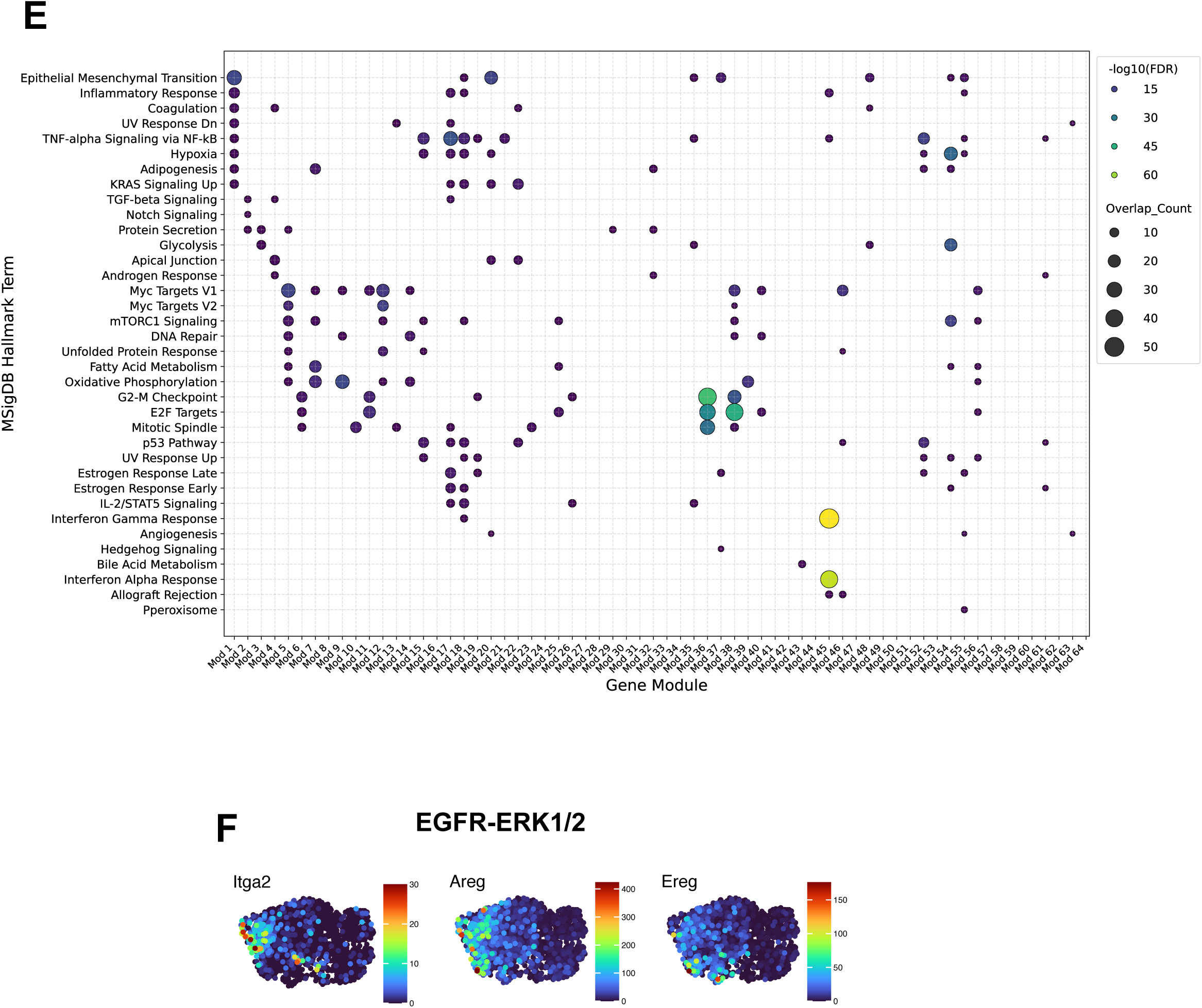
Related to EMT-associated tumor cell heterogeneity emerges along two transcriptional trajectories during C3-TAg tumor progression. **A.** Monocle3 trajectory analysis of tumor cells from 5-mm and >15-mm C3-TAg tumors shown in Fig. 2A-C. Top row shows the raw trajectory analysis. Bottom row shows the overlay of the trunk and branching into EMT trajectory 1 and EMT trajectory 2. **B.** Expression of representative mammary lineage, epithelial, and mesenchymal-associated genes across pseudotime for EMT trajectory 1. **C.** Expression of representative mammary lineage, epithelial, and mesenchymal-associated genes across pseudotime for EMT trajectory 2. **D.** Heatmap showing scaled activity scores for inferred gene modules across tumor cell Seurat clusters, with hierarchical clustering of modules and clusters. Highlighted modules correspond to programs shown in Fig. 2F. **E.** Dot plot showing Hallmark pathway enrichment across inferred gene modules. Dot size indicates the number of overlapping genes, and color indicates enrichment significance. **F.** UMAP feature plots showing expression of selected EGFR–ERK1/2-associated genes, including Itga2, Areg, and Ereg, across tumor cells.

**Figure S3.**
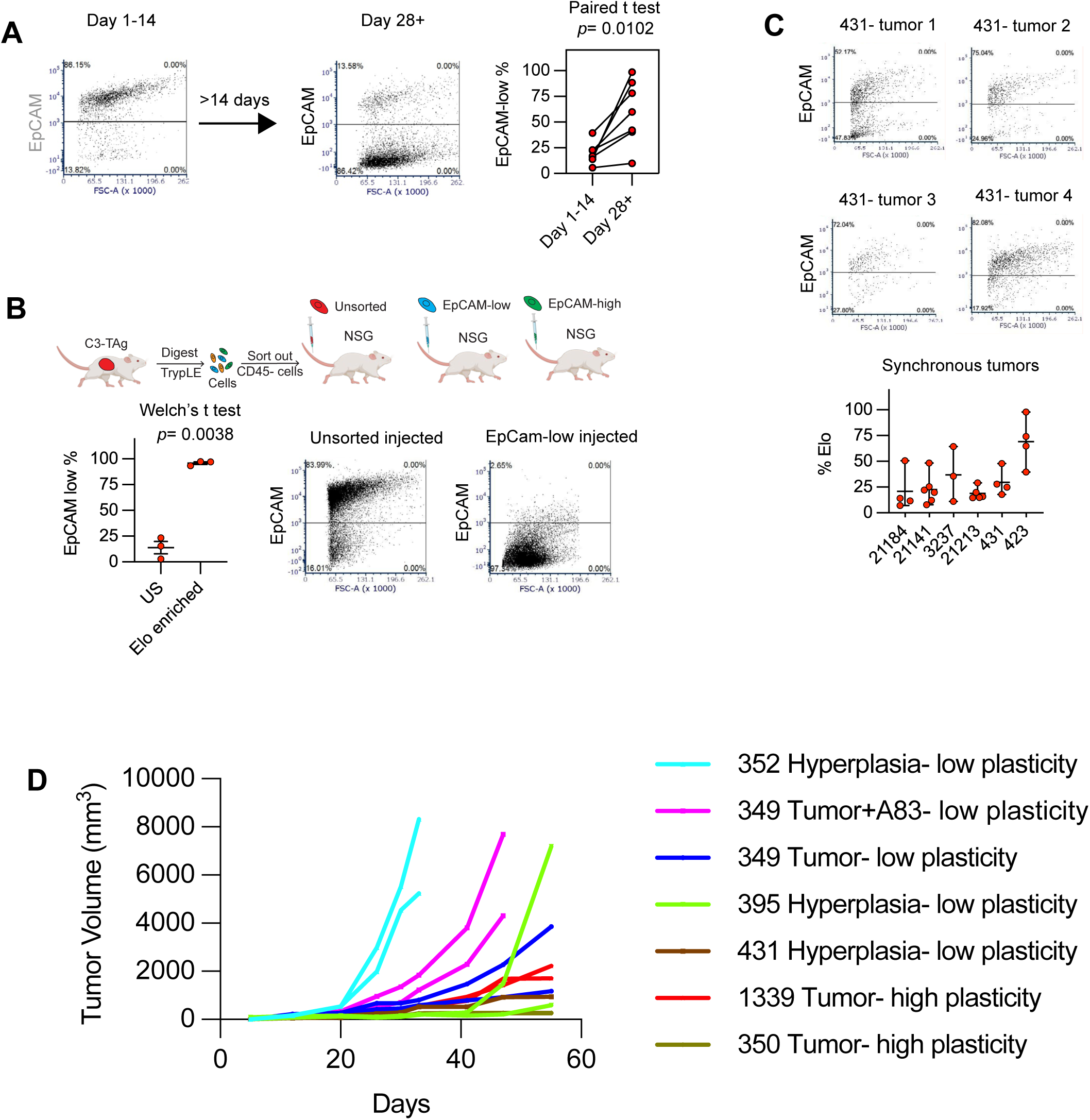
Related to EMT-associated plasticity is acquired during tumor progression and generates stable mesenchymal-like states. **A.** Representative FACS plots and paired quantification showing the percentage of EpCAM-low cells in C3-TAg tumor organoids at early culture time points and after extended culture. **B.** Experimental workflow for isolating CD45-negative unsorted, EpCAM-low, and EpCAM-high C3-TAg tumor cells and transplanting sorted populations into NSG mice. Representative FACS plots and quantification show enrichment of EpCAM-low cells in tumors derived from EpCAM-low injected cells compared with unsorted controls. **C.** Representative FACS plots and quantification showing variability in the percentage of Ep-CAM-low cells across synchronous tumors. **D.** Tumor growth curves for independent hyperplasia- and tumor-derived organoid lines classified as low- or high-plasticity based on ability to convert to an E-lo state.

**Figure S4.**
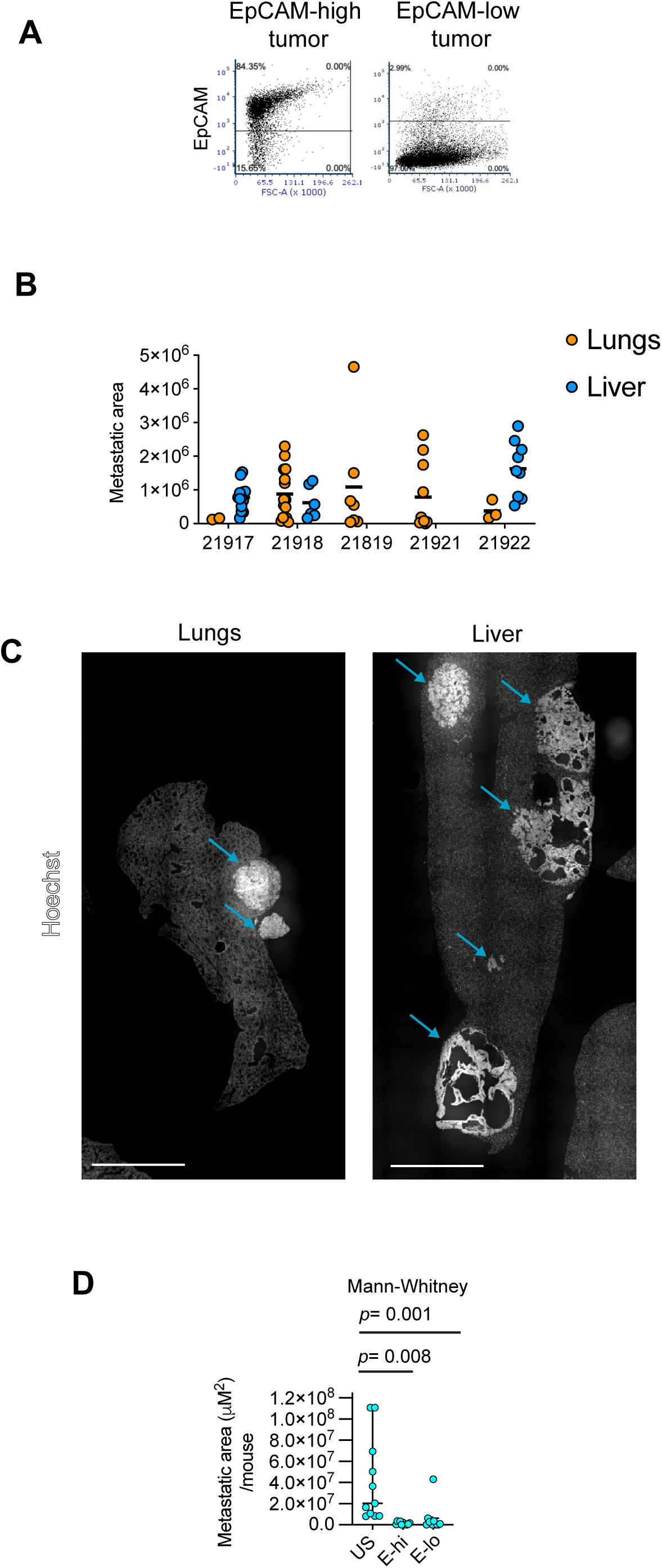
Related to hybrid EpCAM-high and mesenchymal-like EpCAM-low C3-TAg tumor cells are both metastatically competent. **A.** Representative FACS plots showing C3-TAg tumor lines enriched for EpCAM-high or Ep-CAM-low tumor cell states. **B.** Quantification of metastatic lesions in lungs and liver following intracardiac injection of Ep-CAM-high-enriched C3-TAg tumor cells. Each mouse is shown individually, with metastatic area quantified for lung and liver lesions. **C.** Hoechst-stained tissue sections showing metastatic lesions in lungs and liver following intracardiac injection of EpCAM-high-enriched C3-TAg tumor cells. Arrows indicate metastatic lesions. **D.** Expanded quantification from Fig. 4F that includes the Unsorted tumor cell injections. The E-hi and E-lo plots are the same data from Fig. 4F. Quantification of metastatic area per mouse following injection of unsorted, EpCAM-high, or EpCAM-low C3-TAg tumor cells.

**Figure S5.**
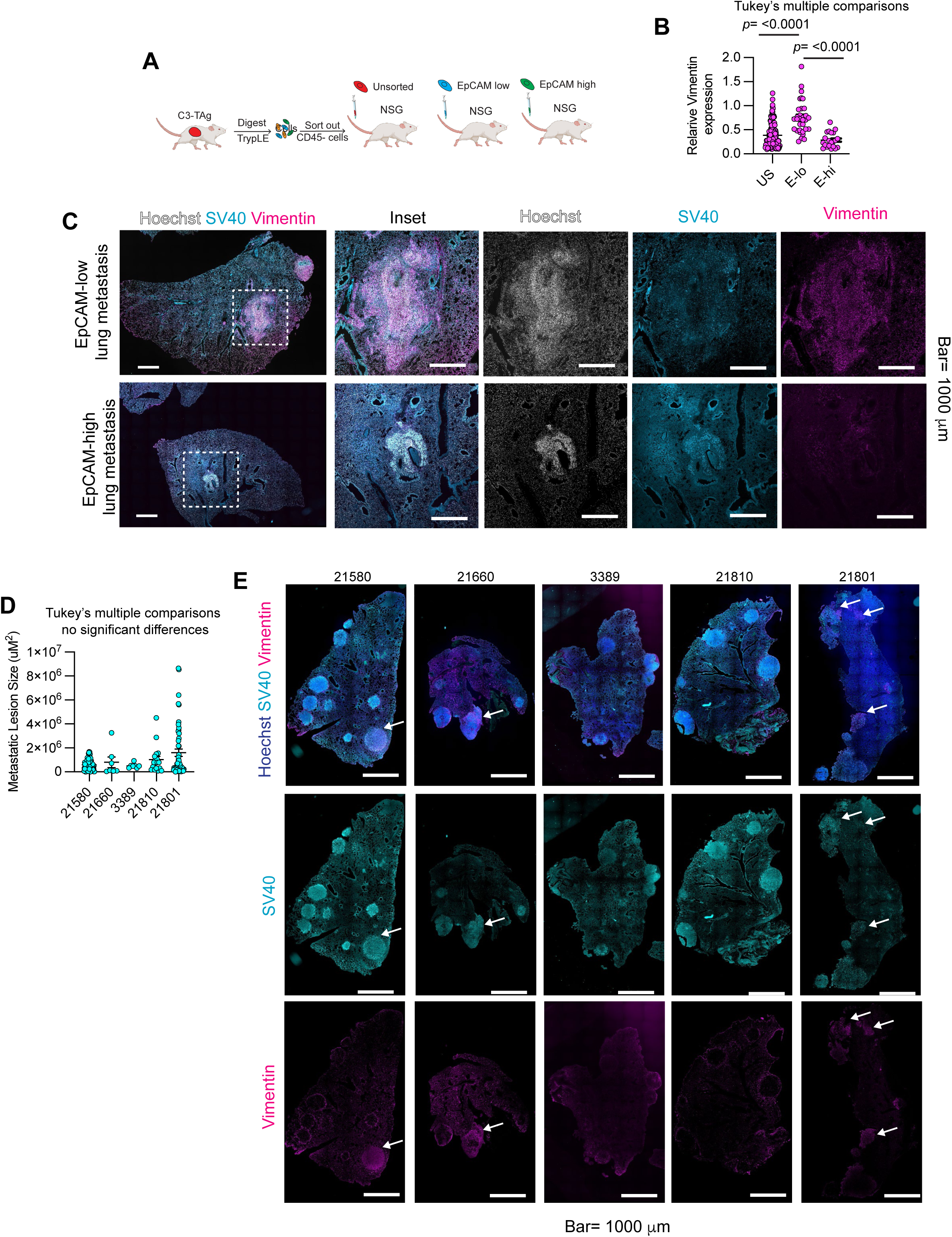
Related to metastatic lesions derived from EpCAM-high and EpCAM-low C3-TAg tumor cells retain distinct EMT-associated phenotypes. **A.** Experimental workflow for direct FACS isolation of CD45-negative unsorted, EpCAM-high, and EpCAM-low C3-TAg tumor cells followed by injection into NSG mice and analysis of lung metastases. **B.** Quantification of relative Vimentin expression in lung metastases generated by unsorted, EpCAM-low, or EpCAM-high tumor cells. **C.** Representative immunofluorescence images of EpCAM-low- and EpCAM-high-derived lung metastases stained for Hoechst, SV40, and Vimentin. Scale bars, 1000 μm. **D.** Quantification of metastatic lesion size across independent C3-TAg tumor lines. No significant differences were detected by one-way ANOVA with Tukey’s multiple-comparisons. **E.** Representative immunofluorescence images of lung metastases generated from independent C3-TAg tumor lines stained for Hoechst, SV40, and Vimentin. Arrows indicate metastatic lesions or regions of Vimentin expression. Scale bars, 1000 μm.

**Table S1. Differentially expressed marker genes for tumor epithelial cell clusters.** Marker genes defining each re-clustered C3-TAg tumor epithelial cell population after removal of canonical cell-cycle genes. For each gene, the table reports the associated Seurat cluster, average log2 fold change, percentage of cells expressing the gene within and outside the cluster, nominal P value, and adjusted P value.

**Table S2. Hallmark pathway enrichment analysis of tumor epithelial cell clusters.** MSigDB Hallmark pathway enrichment analysis performed on cluster-defining genes from re-clustered C3-TAg tumor epithelial cells. The table reports enriched Hallmark terms for each cluster, including cluster size, gene-set overlap, nominal and adjusted P values, and overlapping genes. Significant Hallmark enrichments highlight EMT-associated programs in Clusters 2, 3, 6, and 8.

**Table S3. Gene Ontology enrichment analysis of tumor epithelial cell clusters.** Gene Ontology enrichment analysis of cluster-defining genes from re-clustered C3-TAg tumor epithelial cells. The table lists enriched GO biological process terms for each cluster, including cluster size, gene-set overlap, nominal and adjusted P values, and overlapping genes. These enrichments support functional differences among tumor epithelial states, including programs associated with proliferation, epithelial differentiation, EMT, motility, and extracellular matrix organization.

**Table S4. Gene module assignments for tumor epithelial cells.** Gene module assignments derived from tumor epithelial cell trajectory analysis. The table lists each gene, its assigned module and supermodule, and the corresponding dimensional coordinates used to organize module structure.

**Table S5. Hallmark pathway enrichment analysis of tumor epithelial gene modules.** MSigDB Hallmark pathway enrichment analysis performed on genes assigned to each tumor epithelial cell module. The table reports enriched Hallmark terms for each module, including module size, gene-set overlap, nominal and adjusted P values, and overlapping genes.

**Table S6. Gene Ontology biological process enrichment analysis of tumor epithelial gene modules.** Gene Ontology biological process enrichment analysis performed on genes assigned to each tumor epithelial cell module. The table lists enriched biological process terms for each module, including module size, overlap, nominal and adjusted P values, and overlapping genes.

**Table S7. Tabula Muris cell-type enrichment analysis of tumor epithelial gene modules.** Enrichment analysis comparing tumor epithelial gene modules with Tabula Muris cell-type signatures. The table reports enriched cell-type terms for each module, including module size, gene-set overlap, nominal and adjusted P values, and overlapping genes.

**Table S8. Gene Ontology cellular component enrichment analysis of tumor epithelial gene modules.** Gene Ontology cellular component enrichment analysis performed on tumor epithelial gene modules. The table lists enriched cellular component terms for each module, including module size, overlap, nominal and adjusted P values, and overlapping genes.

**Table S9. Gene Ontology molecular function enrichment analysis of tumor epithelial gene modules.** Gene Ontology molecular function enrichment analysis performed on tumor epithelial gene modules. The table reports enriched molecular function terms for each module, including module size, overlap, nominal and adjusted P values, and overlapping genes.

**Table S10. Enrichment of tumor epithelial gene modules for TGFβ- and EGFR–ERK1/2-regulated programs.** Enrichment analysis comparing tumor epithelial gene modules with gene signatures induced by EGF or TGFβ treatment and genes suppressed by erlotinib or trametinib in C3-TAg tumor organoids. The table includes enrichment scores, nominal and FDR-adjusted P values, overlap counts, and overlapping genes.

**Table S11. Enrichment of tumor epithelial clusters for TGFβ- and EGFR–ERK1/2-regulated programs.** Enrichment analysis comparing tumor epithelial Seurat cluster gene signatures with TGFβ-induced, EGF-induced, erlotinib-suppressed, and trametinib-suppressed gene sets from C3-TAg tumor organoids. The table reports overlap counts, nominal P values, and enrichment significance for each cluster-signature comparison.

